# Microbiome Dynamics of Bovine Mastitis Progression and Genomic Determinants

**DOI:** 10.1101/2020.07.13.200808

**Authors:** M. Nazmul Hoque, Arif Istiaq, M. Shaminur Rahman, M. Rafiul Islam, Azraf Anwar, AMAM Zonaed Siddiki, Munawar Sultana, Keith A. Crandall, M. Anwar Hossain

## Abstract

The milk of lactating cows presents a complex ecosystem of interconnected microbial communities which can impose a significant influence on the pathophysiology of mastitis. Previously, we reported the alteration of microbiome (bacteria, archaea, virus) composition between clinical mastitis (CM) and healthy (H) milk. We hypothesized possible dynamic shifts of microbiome compositions with the progress of different pathological states of mastitis (CM, Recurrent CM; RCM, Subclinical Mastitis; SCM) determined by its favoring genomic potentials. To evaluate this hypothesis, we employed whole metagenome sequencing (WMS) in 20 milk samples (CM = 5, RCM = 6, SCM = 4, H = 5) to unravel the microbiome dynamics, interrelation, and relevant metabolic functions. PathoScope (PS) and MG-RAST (MR) analyses mapped the WMS data to 442 bacterial, 58 archaeal and 48 viral genomes with distinct variation in microbiome composition and abundances across these metagenomes (CM>H>RCM>SCM). PS analysis identified 385, 65, 80 and 144 bacterial strains in CM, RCM, SCM, and H milk, respectively, with an inclusion of 67.19% previously unreported opportunistic strains in mastitis metagenomes. Moreover, MR detected 56, 13, 9 and 46 archaeal, and 40, 24, 11 and 37 viral genera in CM, RCM, SCM and H-milk metagenomes, respectively. The CM-microbiomes had closest association with RCM-microbiomes followed by SCM, and H-microbiomes. Furthermore, we identified 333, 304, 183 and 50 virulence factors-associated genes (VFGs), and 48, 31, 11 and 6 antibiotic resistance genes (AGRs) in CM, RCM, SCM, and H-microbiomes, respectively, showing a significant correlation between the relative abundances of VFGs (p = 0.001), ARGs (p = 0.0001), and associated bacterial taxa. We also detected correlated variations in the presence and abundance of several metabolic functional genes related to bacterial colonization, proliferation, chemotaxis, motility and invasion, oxidative stress, virulence and pathogenicity, phage integration and excision, biofilm-formation, and quorum-sensing to be associated with different episodes of mastitis. Therefore, profiling the dynamics of microbiome in different states of mastitis, concurrent VFGs, ARGs, and genomic functional correlations will contribute to developing microbiome-based diagnostics and therapeutics for bovine mastitis, and carries significant implications on curtailing the economic fallout from this disease.

## Introduction

Mastitis is the inflammation of the mammary gland and/or quarters^1^ and represents the foremost disease-driven challenge in production faced by the global dairy industry^2^. The milk microbiota composition is an important determinant of mammalian health^3, 4^, plays an important role in udder health by interacting with the immune and metabolic functions of the cow^5, 6^, the spread of virulence factors^7^, and the spread of antimicrobial resistance genes (ARGs)^8, 9^. Previously, we reported that the microbiome signature in bovine clinical mastitis (CM) has functional biasness from a genomic standpoint^6^, and observed distinct shifts and differences in abundance between the microbiome of CM and H milk. Furthermore, bovine CM milk had a 68.04% inclusion of previously unreported and/or opportunistic strains^6^. In addition, a plethora of archaeal and viral entities^6^ might also be present in association with the bacteria, suggesting relevant physiological and pathological implications for their host. Although bovine mastitis microbiome comprises of bacteria, archaea, viruses, and other microorganisms, until now, most researchers have mainly focused on the bacterial component of the milk microbiota^11–14^.

We also reported that the resistomes (the total collection of antibiotic resistance genes) act as a potential key factor in disease complication and recurrence, could be concurrent to microbiome signature^10^. However, no association was found with cattle breed, despite significant differences in microbiome diversity among different breeds^10^. Bovine mastitis microbiomes may show a shift in taxonomic diversity and composition according to disease states (clinical, subclinical and recurrent)^6, 11, 14, 15^. Following pathogen invasion and subsequent establishment in the mammary gland, either CM or subclinical mastitis (SCM) may present itself. Bovine CM is one of the most frequent diseases (with visibly abnormal milk, and swollen, hot, painful, and reddened udder) affecting the global dairy industry^16, 17^, and diverse groups of microbial communities colonizing the mammary gland and/or quarters have evolved novel mechanisms that facilitate their proliferation during the disease process^6^. One of the very frustrating aspects of bovine CM is its recurrent nature, reinfection of a quarter or udder after bacteriological cure^18^. Recurrent CM (RCM) is caused by persistent intra-mammary infections (IMI)^17^, and specific differences in the milk microbiota may contribute to recurrence susceptibility^18^. IMI may persist beyond the resolution of the clinical symptoms of CM, and sub-sequent RCM flare-ups may be observed^17^. Almost every dairy herd has cows with SCM^15^, and a variety of pathogens can establish chronic infections which may only occasionally manifest the clinical signs of mastitis^19^.

During the progression of mastitis, dysbiosis in the milk microbiome can occur, with an increase in opportunistic pathogenic bacteria, and a reduction in healthy milk commensal bacteria^6, 20, 21^. Researchers have now started to explore the complexities of the microbial ecology within this niche through disease progression^6, 7, 10, 20^. While understanding the role of all interacting microbial players contributing to disease progression is becoming increasingly crucial in the face of large-scale epidemics^22, 23^, such a study has yet to be completed for dairy animal mastitis. The microbial community present within the mammary gland and/or quarters is being increasingly recognized as a key regulator of metabolic and immune homeostasis^5, 6^, and a mediator of antimicrobial resistance^6^. The resistomes or ARGs, which exist in both pathogenic^9, 10^ and nonpathogenic commensal bacteria^9^ are frequently carried on mobile genetic elements. Similarly, the virulome, the set of genes encoding virulence, can also be carried on the mobilome^9^, and thus, many virulence factors associated genes (VFGs) easily spread in bacterial populations by horizontal gene transfer, converting mutualistic or commensal bacteria into potential opportunistic pathogens^7, 21, 24^. Though the exact functions of archaea and viruses in the progression of mastitis are not fully explored like bacterial pathogens^6, 10, 16^, the dynamic shifts in their composition and relative abundances in the onset of CM, persistent or chronic recurrent CM (RCM), invisible udder infections (SCM), and the healthy conditions of the mammary glands and/or quarters could be resulted from a host genetic predisposition to colonization, immune responses to milk bacteria involving increased levels of polymorphonuclear infiltrate, differences mediated by the microbial composition itself, or a combination of these effectors^6, 12, 24^. However, it’s not fully understood what exactly happens to the microbiome when disease states change; which microbiomes are responsible, and what properties enable them to persist through different disease states. Consequently, treatment of these conditions could be exceedingly difficult rendering ineffectiveness of the therapeutics to alleviate the disease. Therefore, we adopted a comprehensive study encompassing all the mastitis categories and investigated their dynamics with genomic properties.

We employed the high-throughput WMS technique (on an average 20.83 million reads per sample, Supplementary Data 1) to gain insight into the phylogenetic composition and diversity of these microbiomes^6, 26^, and to profile their functional attributes^6, 27^ in 15 mastitis (CM, RCM, SCM) and 5 healthy (H) milk samples. We investigated whether consistent and specific disease-associated changes in the microbial communities and their genomic properties could be identified across multiple sample categories of the same disease, and in the healthy condition, of the mammary glands. Culture-independent WMS investigation of the bovine milk microbiome suggested the presence of diverse microbes in milk from healthy and infected mammary glands far greater than previously described^6, 12, 25^.

Moreover, the overexpression of genes related to ARGs, VFGs, and several metabolic pathways coming from the metabolic activities of the microbiome is a crucial factor for the development and progression of bovine mastitis^6^. Therefore, our study aims to describe, for the first time, a comprehensive scenario of the alterations in microbiome composition along with their VFGs, ARGs, and metabolic potentials in different pathological categories of bovine mastitis (CM, RCM, SCM), and healthy milk through the WMS approach.

## Results

The WMS of 20 milk (CM = 5, RCM = 6, SCM = 4, H = 5) samples generated 416.65 million raw reads with an average of 20.83 million reads per sample (range 2.72–39.75 million) (Supplementary Fig. 1, Supplementary Data 1). A total of 385.53 million reads, averaging 19.28 million reads per sample, passed quality control by BBDuk^6^, and were simultaneously analyzed using two different bioinformatics tools, PathoScope 2.0 (PS)^28^, and MG-RAST (MR)^29^. This WMS study produced a direct and comprehensive evaluation of the microbiome diversity and composition, VFGs, ARGs, and functional metabolic potentials of microbiomes to be associated with mammary gland pathogenesis in lactating cows.

### Mastitis-associated changes in the milk microbial diversity and community structure

To evaluate the effect of bovine mastitis categories (CM, RCM, SCM) on the milk microbial alpha diversity (i.e., within-sample diversity), we computed the number of species and/or strains observed in each sample (i.e., richness), and Chao1, ACE, Shannon, Simpson and Fisher indices (i.e., a diversity index accounting both evenness and richness) at the strain level. We found that both the number of observed species/strains, Chao1, ACE, Shannon, Simpson and Fisher indices estimated diversity significantly varied across the four sample groups (CM, RCM, SCM and H) (p = 0.003, Kruskal-Wallis test), and were higher in CM and RCM groups compared to the groups of SCM and H-milk samples (Fig. 1a,b). The rarefaction curves reached a plateau after, on average, 19.28 million reads (Supplementary Fig. 1, Supplementary Data 1)- suggesting that the depth of coverage was sufficient to capture the entire microbial diversity within each sample. Principal coordinate analysis (PCoA), as measured on the Bray-Curtis distance method (Fig. 1c), and non-metric multidimensional scaling (NMDS) ordination plots, through weighted-UniFrac distance method (Fig. 1d), showed discrimination across the metagenomes. The CM-microbiomes clustered more closely to the RCM-microbiomes followed by SCM-microbiomes, and more distantly clustered with H-milk microbes (Fig. 1c, d). Therefore, significant variation in microbiome diversity and composition across these four categories (p = 0.001, Kruskal-Wallis test, Beta diversity test Fig.1 c,d) was also evident. Furthermore, by annotating the WMS reads against different reference databases (namely, virulence factor database (VFDB)^30^, ResFinder^31^, and KEGG and SEED modules of MR^29, 32^), we demonstrated that there were distinct differences in mastitis-specific factors such as VFGs, ARGs, and metabolic functions (Supplementary Data 2) in four metagenomes, which could strongly modulate microbiomes dysbiosis, and the pathophysiology of bovine mastitis.

The composition of bovine milk microbiomes at the domain level was numerically dominated by bacteria, with a relative abundance of 99.58%, followed by archaea (0.24%), and viruses (0.18%) (Supplementary Data 1). In this study, the taxonomic abundances of the microbiomes always remained higher in mastitis (CM, RCM and SCM)-metagenomes compared to the H-milk metagenome. However, the taxonomic abundances fluctuated more within mastitis groups. The unique and shared distribution of microbiomes found in the mastitis and healthy-milk samples are represented by comprehensive Venn diagrams (Fig. 2). We detected 18 bacterial phyla, of which 16, 11, 4 and 13 were found in CM, RCM, SCM and H milk metagenomes, respectively (Supplementary Data 1). Moreover, four bacterial phyla (*Proteobacteria*, *Firmicutes*, *Actinobacteria* and *Bacteroidetes*) were shared across the four groups (Supplementary Fig. 2). The CM milk metagenome had an exclusively unique association with five bacterial phyla (*Deinococcus*, *Acidobacteria*, *Chrysiogenetes*, *Deferribacteres* and *Chloroflexi*), and two typical bacterial phyla (*Thermotogae* and *Verrucomicrobia*) were uniquely present in H-milk samples (Supplementary Data 1). Moreover, 78 orders of bacteria were identified in four metagenomes, of which 13.89%, 4.17% and 6.56% orders were exclusively unique in CM, RCM, and H-metagenomes, respectively (Supplementary Fig. 3, Supplementary Data 1). The MR pipeline simultaneously detected bacteria, archaea, and viruses in these metagenomes. A total of 373 bacterial genera, including 314, 187, 66 and 272 in CM, RCM, SCM and H milk samples, respectively, were detected (Fig. 2a, Supplementary Data 1), and 18.69%, 3.63%, 4.54%, and 9.75% detected genera had sole associations with CM, RCM, SCM and H conditions of the mammary glands. However, 12.06% bacterial genera (Fig. 2a) were common in all of four sample groups (Supplementary Data 1). We detected 442 bacterial strains through PS analysis, of which 385, 65, 80 and 144 strains were found in CM, RCM, SCM and H milk, respectively (Fig. 2b). Of the identified strains, 15.44%, 12.30%, 3.75% and 20.14% strains in CM, RCM, SCM, and H milk, respectively had exclusive associations, and only 4.75% were found to be shared across these metagenome (mastitis and healthy) samples (Fig. 2b). The CM milk microbiome however had 13.1%, 14.8% and 27.6% shared bacterial strains with RCM, SCM and H microbiomes, respectively (Fig. 2b, Supplementary Data 1).

The MR pipeline detected 58 archaeal, and 48 viral genera in four metagenomes, and of them, 12.06% archaeal (Fig. 2c), and 20.83% viral (Fig. 2d) were common in four metagenomes. By comparing these genera across the sample categories, we identified 56, 13, 9 and 46 archaeal, and 40, 24, 11 and 37 viral genera in CM, RCM, SCM and H-milk metagenome, respectively. We found that 23.2%, 16.1% and 75.9% archaeal (Fig. 2c, Supplementary Data 1), and 60.0%, 27.5% and 45.3% viral genera in RCM, SCM and H samples, respectively shared with those of CM samples (Fig. 2d Supplementary Data 1). The CM milk metagenome had sole association of 12 (21.43%) archaeal, and 11 (27.50%) viral genera. The composition of microbial taxa remained much higher in CM milk followed by H, RCM, and SCM-milk metagenomes (Fig. 2e). Despite having lower taxonomic composition in RCM and SCM-metagenomes, the relative abundances of the associated microbiomes varied significantly (p=0.027, Kruskal-Wallis test) across these metagenomes (Supplementary Data 1).

### Mastitis-associated shifts in bacteriome composition

We found that both the composition and the relative abundances of bacterial taxa at the phylum, order, genus, and strain-level differed between mastitis and healthy controls irrespective of the bioinformatics tools used. The associated bacterial phyla were numerically dominated by *Proteobacteria*, *Firmicutes*, *Actinobacteria* and *Bacteroidetes* (comprising > 99% of relative abundance) in mastitis and healthy-milk metagenomes. Additionally, *Proteobacteria* was found to be the single most dominating phylum in CM, RCM and H milk samples (> 90% relative abundance), while in the SCM-milk metagenome, the most abundant bacterial phylum was *Firmicutes* (42.95%) followed by *Bacteroidetes* (33.95%), *Proteobacteria* (22.15%), and *Actinobacteria* (0.95%) (Supplementary Data 1). *Pseudomonadales* was the predominantly abundant bacterial order in CM (86.25%), RCM (90.93%) and H (98.51%) milk whereas *Lactobacillales* (41.59%) and *Flavobacteriales* (31.01%) were the most abundant orders in SCM-milk (Supplementary Fig. 4, Supplementary Data 1). The relative abundance of *Enterobacteriales* also varied among the sample categories and remained significantly higher in the mastitis milk (CM, 12.81%; RCM, 5.42%; SCM, 12.11%) compared to the H milk (0.23%) samples. Although the SCM milk had the lowest number of bacterial orders, their relative abundances remained higher than those detected in the other (CM, RCM, and H) metagenomes (Supplementary Fig. 2, Supplementary Data 1).

We also found significant differences (p = 0.001, Kruskal-Wallis test) in the structure and relative abundance of the bacteria at the genus level. In CM and RCM-metagenomes, *Acinetobacter* was the most abundant bacterial pathogen with a relative abundance of 68.59% and 72.61%, respectively, but remained much lower (0.81%) in SCM-milk samples. The CM milk metagenome was also predominated by *Pseudomonas* (17.69%), *Escherichia* (8.62%), *Shigella* (1.12%), *Klebsiella* (0.66%), *Salmonella* (0.52%), *Enterobacter* (0.50%), *Shewanella* (0.46%) *Pantoea* (0.46%), and *Citrobacter* (0.32%), and rest of the genera had < 0.1% relative abundance (Fig. 3, Supplementary Data 1). Simultaneously, *Pseudomonas* (18.21%), *Aeromonas* (2.64%), *Klebsiella* (1.68%), *Enterobacter* (0.79%), *Escherichia* (0.68%), *Pantoea* (0.61%), *Citrobacter* (0.54%), *Salmonella* (0.52%), *Shewanella* (0.35%), *Lactococcus* (0.24%), and *Cronobacter* (0.13%) were the most abundant genera in the RCM-metagenome (Fig. 3, Supplementary Data 1). Likewise, top abundant bacterial genera in SCM-metagenome were *Lactococcus* (39.20%), *Chryseobacterium* (23.55%), *Ralstonia* (6.65%), *Serratia* (4.43%), *Escherichia* (4.98%), *Riemerella* (2.58%), *Pseudomonas* (1.67), *Citrobacter* (1.40%), *Streptococcus* (1.13%), *Pedobacter* (1.08%) (Fig. 3, Supplementary Data 1). Conversely, the H-milk metagenome was mostly dominated by two bacterial genera, *Acinetobacter* (73.66%) and *Pseudomonas* (24.01%), with relatively lower abundances of *Psychrobacter* (0.62%), *Streptococcus* (0.25%), *Lactococcus* (0.22%), *Ralstonia* (0.18%) (Fig. 3, Supplementary Data 1). Despite having substantially lower (< 0.5) relative abundances, 60, 7, 3 and 27 exclusively associated bacterial genera were found in CM, RCM, SCM and H-milk metagenomes (Supplementary Data 1). These findings suggest that mastitis-associated metagenomes have higher abundance in associated bacterial genera compared to H milk metagenomes, with the inclusion of new and/or the same genera of opportunistic bacteria. However, the relative abundances of mastitis causing bacteria remain more inconsistent according to the different categories of mastitis across CM, RCM and SCM (Supplementary Data 1).

We further investigated the strain-level differences of microbial communities across these four metagenomes through PS analysis which revealed significant variation (p = 0.002, Kruskal-Wallis test) in microbiome composition, diversity and relative abundances across four metagenomes (Fig. 4, Supplementary Data 1). The strain-level microbiome shift was also evident within three mastitis states, with a significantly (p = 0.001, Kruskal-Wallis test) higher number of bacterial strains in the CM-metagenome over the RCM and SCM-metagenomes. Of the detected strains, 62.33%, 13.85% and 17.50% had a unique or sole association with bovine CM, RCM and SCM, respectively (Fig. 2a, Supplementary Data 1). However, the presence of few predominating bacterial species in the four metagenomes (particularly in RCM and SCM) suggests that the crucial differences might also be occurring at the strain-level, and most of the species identified in each sample of the corresponding metagenome were represented by a single strain. The most prevalent bacterial strain, *Acinetobacter johnsonii* XBB1 had almost 1.7-fold higher relative abundance in CM (59.92%) milk over H (36.2%) milk but remained much lower (< 0.5%) in the other two metagenomes (RCM, SCM) (Fig. 4, Supplementary Data 1). Moreover, *Pseudomonas putida* KT2440 (11.81%), *Escherichia coli* O104:H4 str. 2011C (0.036%), *Aeromonas veronii* B565 (1.79%), *Pantoea dispersa* EGD-AAK13 (1.74%), *Klebsiella oxytoca* KA-2 (1.09%), *P. entomophila* L48 (0.80%), *Kluyvera ascorbata* ATCC 33433 (0.68%), *P. japonica* NBRC 103040 (0.64%), *Aeromonas hydrophila* YL17 (0.61%) and *A. tandoii* DSM 14970 (0.56%) were the predominant strains found in CM milk (Fig. 4, Supplementary Data 1). *Micromonospora* sp. HK10 (54.11%), *Campylobacter mucosalis* CCUG 21559 (26.52%), *Catenibacterium mitsuokai* DSM 15897 (11.15%), and *Anaerobutyricum hallii* DSM 3353 (7.09%) were the predominant strains in the RCM-metagenome. The most abundant strains in the SCM-metagenome were *Micromonospora* sp. HK10 (38.29%), *Campylobacter mucosalis* CCUG 21559 (26.25%), *Anaerobutyricum hallii* DSM 3353 (25.74%), and *Catenibacterium mitsuokai* DSM 15897 (8.62%). In contrast, the H-milk metagenome was mostly dominated by *Acinetobacter johnsonii* XBB1 (36.20%), *Micromonospora* sp. HK10 (15.16%), *Anaerobutyricum hallii* DSM 3353 (11.60%), *Campylobacter mucosalis* CCUG 21559 (7.42%), and different strains of *Pseudomonas* such as *P. fragi* str. A22 (16.04%), *P. fluorescens* SBW25 (2.43%), *P. protegens* CHA0 (1.02%), *P. fluorescens* SBW25 (0.70%), *P. alkylphenolica* KL28 (0.65%), and *P. resinovorans* NBRC 106553 (0.52%). The rest of the bacterial strains identified in these metagenomes had much lower (< 0.50%) relative abundances (Fig. 4, Supplementary Data 1).

Across the three mastitis metagenomes, 67.19% of the detected bacterial strains were previously unreported (of which 70.13% were in CM, 44.62% were in RCM and 43.75% were in SCM), and possibly played an opportunistic role in mammary gland pathogenesis. The top abundant opportunistic strains in CM-milk metagenome were *Aeromonas veronii* B565 (19.16%), *Pantoea dispersa* EGD-AAK13 (18.65%), *Klebsiella oxytoca* KA-2 (11.70%), *Kluyvera ascorbata* ATCC 33433 (7.27%), *Aeromonas hydrophila* YL17 (6.54%). In RCM-milk, the most abundant strains were *Nocardia pseudobrasiliensis* (48.06%), *Serratia marcescens* subsp. marcescens Db11 (16.22%), *Nocardia mikamii* NBRC 108933 (15.19%), *Corynebacterium bovis* DSM 20582 (4.66%), and *Aeromonas hydrophila* SSU (3.11%). Different strains of the *Chryseobacterium* genus such as *Chryseobacterium* sp. Leaf405 (19.06%), *Chryseobacterium* sp. CF299 (13.68%), *Chryseobacterium haifense* DSM 19056 (13.19), *Chryseobacterium greenlandense* (13.04%), and *Chryseobacterium* sp. YR460 (12.56%), and *Serratia marcescens* subsp. marcescens Db11 (8.30%) and *Citrobacter freundii* CFNIH1 (6.65%) were the most abundant opportunistic strains in SCM-metagenome (Supplementary Table 2

### Mastitis-associated shift in archaeal and viral fraction of the microbiome

Another noteworthy feature of the current study is the concomitant detection of archaeal and viral fractions of microbiomes in both mastitis and H-milk metagenomes through MR analysis. Interestingly, the structure and relative abundance of several microbial clades of these two domains were decreased in both RCM (11 genera) and SCM (9 genera) milk-metagenomes and remained enriched in CM-milk (56 genera) and healthy controls (46 genera) (Figs. 4,5, Supplementary Data 1). Among the identified archaeal components of the microbiome, *Methanosarcina* was the most abundant genus both in CM (41.78%) and RCM (34.04%) milk samples. Besides this genus, *Methanococcoides* (21.25%), *Methanocaldococcus* (2.43%), *Thermococcus* (1.59%), *Haloterrigena* (0.95%), *Sulfolobus* (0.80%), *Methanothermococcus* (0.71%) and *Methanocorpusculum* (0.59%) in CM-milk, *Methanobrevibacter* (25.53%), *Halogeometricum* (10.64%) and *Pyrococcus* (6.38) in RCM-milk, and *Methanococcus* (24.39%), *Methanosphaera* (19.51%), *Methanoregula* (7.31%), *Methanospirillum* and *Methanothermobacter* (4.88% each) in SCM-milk metagenomes were the predominantly abundant archaeal genera (Fig. 5, Supplementary Data 1). On the other hand, *Haloarcula* (12.94%), *Methanoplanus* (11.95%), *Euryarchaeota* (4.12%), *Methanoculleus* (3.84%), *Methanosaeta* (1.28%), *Archaeoglobus* (1.0%), *Natrialba* and *Halogeometricum* (1.13%, each) were the most abundant archaeal genera in H-milk metagenome (Fig. 5, Supplementary Data 1). Remarkably, the 81.03% and 77.59% of the detected archaeal genera were absent in RCM and SCM-metagenomes, respectively (Supplementary Data 1). The relative abundance of *Haloquadratum* and *Natronomonas* genera remained more than two-fold higher in SCM and RCM samples than CM and H milk samples (Fig. 5). The relative abundance of the rest of the genera remained much lower (< 0.05%) but varied significantly across the four metagenomes (Fig. 5, Supplementary Data 1). Moreover, 13 (23.21%) and 2 (4.35%) archaeal genera in CM and H milk metagenomes, respectively, had unique associations while none were found to be unique in RCM and SCM milk metagenomes. Among these unique genera, *Acidilobus*, *Aciduliprofundum*, *Candidatus*, *Pyrobaculum*, *Halobacterium*, *Thermosphaera* etc. were found in CM while only *Ferroglobus* and *Methanopyrus* were identified in H-milk samples (Supplementary Data 1).

The viral fraction of the microbiome was largely dominated by the members of the *Ortervirales* and *Caudovirales* orders (> 90.0% of the total abundance), represented mostly by *Retroviridae* (52.84%), *Siphoviridae* (21.87%), *Podoviridae* (12.96%), *Myoviridae* (11.21%); the remaining families had relatively lower abundances (< 0.5%) (Supplementary Data 1). The CM milk metagenome had the highest number (40) of viral genera followed by H (37), RCM (24) and SCM (11) milk metagenomes, and of them, only 27.50% and 35.13% viral genera had sole association in CM and H-milk metagenomes, respectively. Besides, 60.0%, 27.50% and 45.3% genera in RCM, SCM and H-milk metagenomes, respectively were found to be shared with CM-milk metagenome (Supplementary Data 1). In this study, *Siphovirus* was the predominantly identified viral genus among the four metagenomes, but the relative abundance of this genus remained much higher in mastitis (SCM = 50.42%, RCM = 24.76%, CM = 20.33%) milk compared to H (13.35%) milk samples (Fig. 6, Supplementary Data 1). In addition to *Siphovirus*, the most abundant viral genera in CM-metagenome were *P2-like viruses* (13.67%), *Podovirus* (11.57%), *Myovirus* (7.78%) *Bpp-1-like viruses* (6.60%), phiKZ-like viruses (4.06%) and *T1-like viruses* (0.99%). Similarly, *Epsilon15-like viruses* (23.78%), *P1-like viruses* (2.53%), *P22-like viruses* (1.85%), *T4-like viruses* (1.75%), *Caudovirus* (0.97%) and *N15-like viruses* (0.88%) in RCM-milk, and *Lambda-like viruses* (10.08%), *Inovirus* (5.88%), *Molluscipoxvirus* (5.88%) and *phiKZ-like viruses* (4.20%) in SCM-milk metagenomes were the predominant viral genera (Fig. 6, Supplementary Data 1). The H-milk metagenome, however, had a relatively higher abundance of *Betaretrovirus* (46.42%) and *Gammaretrovirus* (30.07%) genera. The rest of the viral genera detected in both mastitis and H-milk metagenomes had much lower (< 0.5%) relative abundances but their abundances always remained higher in CM metagenome (Fig. 6, Supplementary Data 1). Despite having lower relative abundances (< 0.5%), *Epsilonpapillomavirus*, *Gammapapillomavirus*, *Lymphocryptovirus*, *Mastadenovirus*, *Microvirus* etc. in CM-milk, and *AHJD*-like viruses, *Alpharetrovirus*, *Avipoxvirus* c2-like viruses, *Cervidpoxvirus*, *Epsilonretrovirus*, *Lentivirus*, *Parapoxvirus*, *Retrovirus* in H-milk metagenomes were found as the unique viral genera (Supplementary Data 1).

### Virulence factors gene families in mastitis and healthy milk metagenomes

We next sought to identify VFGs among the microbiomes of the four metagenomes through mapping the WMS reads against the virulence factor database (VFDB)^30^. The VFDB annotations revealed a clear, and distinct enrichment of the predicted VFGs in association with bovine mastitis. We detected 494 VFGs which showed significant (p = 0.001, Kruskal-Wallis test) differences both in composition, and relative abundances among these four sample categories (Supplementary Data 2). The CM-associated microbiomes had the highest number of VFGs (333), followed by RCM (304), SCM (183) and H (50) milk microbes, and only 3.24% of the detected VFGs were shared among the microbiomes of the four metagenomes (Fig. 7a,b). In this study, 77.4%, 10.3% and 13.3% of the detected VFGs in RCM, SCM and H-milk microbiota, respectively, had a shared association with CM-microbiomes (Fig. 7a, Supplementary Data 2). The virulence properties (number and relative abundances of the VFGs) of the microbiomes were strongly correlated (p = 0.001, Kruskal-Wallis test) to their relative abundances (at the genus level) in the corresponding CM, RCM, SCM, and H-milk metagenomes (Supplementary Table 3). By comparing the relative abundance of the detected VFGs among the microbiomes of four metagenomes, we found that innate immune response associated capsular genes (*ABZJ_00085* and *ABZJ_00086*) of *Acinetobacter* remained higher in H-milk microbiomes (10.05% and 6.87%, respectively) followed by CM (5.58% and 5.26%, respectively), and RCM (4.07% and 3.55%, respectively)-causing microbes, corroborating the higher taxonomic abundance of this genus in H-milk. Surprisingly, these two genes were absent in SCM-milk microbiomes (Fig. 7b, Supplementary Data 2). Importantly, the CM and RCM-microbiomes had sole association of several VFGs including genes encoding crucial nutrient (iron) uptake and/or regulation, *pvd*L (5.11% and 5.28%, respectively), twitching motility and/or chemotaxis, *chp*A (3.17% and 2.90%, respectively) and *mot*C (0.55% and 0.44%, respectively), bacterial adhesion, invasion, or intracellular survival, *mot*B/*omp*A (2.1% and 2.34%, respectively), biofilm-formation, DNA uptake, adhesion to host cells and adherence, *pil*J (1.83% and 1.81%, respectively) and *pvd*A (0.56% and 0.60%, respectively), two-component system, *gac*S (1.55% and 1.45%, respectively),, intracellular multiplication factor, *icm*F1 (1.51% in each), master regulator of biofilm initiation, *bfm*R (1.57% and 1.24%, respectively), quorum-sensing/antibiotics susceptibility altering, *pvd*Q (0.88% and 0.96%, respectively), cellular adherence, *pil*C (0.88% and 0.91%, respectively), heat shock or stress-related, *muc*D (0.65% and 0.55 %, respectively), bacterial communication, *hdt*S (0.65% and 0.55%, respectively), and bacterial pathogenesis, *crc* (0.54% and 0.24%, respectively) (Fig. 7b, Supplementary Data 2). The SCM-microbiomes possessed a higher number of genes coding for bacteriocin production, and cytotoxic activity in *E. coli*, *EcSMS35_B0007* (5.28%), capsular proteins in pathogenic *Klebsiella* species, *cps*B (2.60%) and *rfb* (1.77%), fimbrial proteins, *stg*C (2.10%), and adhesin proteins, *upa*G/*eha*G (2.09%) in pathogenic *E. coli* (Fig. 7b, Supplementary Data 2). Conversely, genes coding for bacterial chemotaxis, *omp*A (3.43%) and *mot*A (1.63%), flagellar assembly, *fli*M (3.08%), *flh*A (2.90%), *flg*I (2.18%), *fli*G (2.13%), *flg*G (1.87%), *fli*F (1.86%), *fli*A (1.47%) and *fli*P (1.41%), biofilm-formation, *alg*8 (3.93%), *alg*I (2.82%), and *alg*D (2.76%), alginate biosynthesis, *alg*G (4.28%) and *alg*U (1.62%), pili and flagella expression/adherence, *rpo*N (4.87%), *fle*Q (3.30%) and *fle*N (1.69%), and outer membrane proteins, *opr*F (3.33%) had relatively over expression in H-milk microbiomes compared to mastitis-causing microbes (Fig. 7b). Although, the rest of the VFGs detected in this study had relatively lower abundances (< 0.5%), they remained predominantly identified in both the CM and RCM-associated microbiomes (Supplementary Data 2).

### Antibiotics resistance gene families in mastitis and healthy milk metagenomes

We further investigated the total number, and classes of different antibiotic resistance genes (ARGs) present in the microbial genomes of these four metagenomes using ResFinder^31^. In our current WMS data set, there was a broad variation in ARGs diversity and composition (Fig. 8a, Supplementary Data 2). The categories and relative abundances of the ARGs were significantly correlated (p = 0.0001, Kruskal-Wallis test) with the relative abundance of the associated bacteria found in CM, RCM, SCM, and H-milk metagenomes (Supplementary Data 2). ResFinder identified 58 ARGs belonged to 18 antibiotic classes distributed in 442 bacterial genomes (Supplementary Data 2). The CM-associated microbiomes harbored the highest number of ARGs (48), followed by RCM (31), SCM (11) and H (6) milk microbes (Fig. 8a). However, the RCM-causing bacteria possessed the highest number (12) of ARG classes followed by CM (11), SCM (9) and H (4) milk bacteria (Supplementary Data 2). In addition, 36.2%, 20.4% and 12.5% of the ARGs in RCM, SCM and H-milk microbes, respectively had shared association with those of CM-pathogens (Fig. 8b). Of the resistant classes, beta-lactams (11 genes) were detected as the most prevalent antibiotic group in all metagenomes followed by tetracyclines (5 genes), quinolones (5 genes), macrolides (10 genes), and trimethoprim (3 genes) (Fig. 8b, Supplementary Data 2). The relative abundances of these ARGs varied across the four metagenomes (Fig. 8 b,c). The broad-spectrum beta-lactams resistant gene, *bla*_OXA_ was found as the common ARG among the microbiomes of four metagenomes, displaying the highest relative abundance (97.63%) in H-milk microbiota followed by RCM (46.17%), SCM (42.54%) and CM (40.80%) microbiomes (Fig. 8b,d, Supplementary Data 2). However, other beta-lactams resistant genes such as *bla*_ROB_ in CM (3.73%) and SCM (1.27%), *bla*_PLA1a_ in CM (0.31%), RCM (0.29%) and SCM (1.16%), and *bla*_CARB-2_ in RCM (0.2%) causing pathogens were also identified (Fig. 8d). The tetracyclines (doxycycline and tetracycline) resistant gene, *tet* had the highest relative abundance (37.62%) in SCM-associated bacteria compared to RCM (25.84%) and CM (13.33%)–pathogens, and was not detected in the H-milk microbiomes (Fig. 8b). Another tetracycline resistant gene, *tet*A was only found in the RCM milk-microbiome with a relative abundance of 1.39% (Fig. 8b). Aminoglycoside (*aph*) and sulfonamide (*sul*) resistant genes were ubiquitously prevalent among the microbes of the four metagenomes, but their relative abundances remained several-folds (> 5) higher in CM-pathogens compared to RCM, SCM and H milk. Macrolides (erythromycin and streptogramin B) resistant (*msr*E), and fosfomycin resistant (*fos*A) genes had higher relative abundances (2.73% and 1.15%, respectively) in RCM- pathogens compared to other mastitis causing bacteria. The multidrug-resistant gene, *oqx*B had the highest relative abundance in RCM (15.11%) while *mdf*A remained predominantly abundant in CM (3.31%) pathogens (Fig. 8b). Interestingly, the *dfr*A (trimethoprim resistant, 2.01%) and *flo*R (florfenicol resistant, 0.24%) genes were identified in only the CM-associated microbial genomes (Fig. 8b). Similarly, the RCM-microbiome also had a unique association with four ARGs; the *cml*A (chloramphenicol resistant), *aad*A (aminoglycoside resistant), *bla*_CARB-2_ (beta-lactam resistant), and *qnr*S (quinolone resistant) genes which were absent among the microbes of other metagenomes (Fig. 8b, Supplementary Data 3). In general, beta-lactam resistant genes (*bla*_OXA_ particularly) presented the highest level of expression (Fig. 8c.d) followed by tetracycline (*tet* and *tet*A), quinolone (*oqx*B, *mdf*A and *qnr*S), and macrolide (*mph*E, *msr*E and *mdf*A) resistant genes. The rest of the ARGs also varied in their expression levels across the four metagenomes, being more prevalent in the RCM-microbiomes (Fig. 8c). By investigating the possible mechanisms of the detected ARGs, we found that enzymatic inactivation associated resistance had the highest level of expression in the microbiomes of the corresponding metagenomes followed by antibiotic efflux pumps, antibiotic inactivation, enzymatic modification, antibiotic target protection/alteration, and folate pathway antagonist-attributed resistance mechanisms (Fig. 8e, Supplementary Data 2).

### Mastitis-associated changes in metabolic functional potentials of the microbiome

The functional annotation of the WMS reads identified 8.55% and 5.98% putative genes with known and unknown protein functions, respectively (Supplementary Data 1). By examining the correlation between the different gene families of the same KEGG pathway for mastitis and H-milk microbiomes, we found significant differences (p = 0.002, Kruskal-Wallis test) in their relative abundances, and positive correlation with different states of bovine mastitis. We found that genes coding for citrate metabolism (TCA cycle) had a more than two-fold overexpression in the CM (16.55%) and SCM (12.50%)-associated microbiomes compared to RCM (6.75%) and H (8.79%)-milk microbiota. In addition, genes encoding citrate synthase (*glt*A) had higher relative abundance in the RCM-microbiome (31.25%), followed by SCM (21.57%), CM (15.41%) and H-milk microbes (13.86%) (Fig. 9a, Supplementary Data 2). The RCM and SCM- milk microbiomes had higher relative abundances of genes associated with oxidative phosphorylation; OP (74.04% and 78.86%, respectively) and carbon fixation pathways in prokaryotes (4.81% and 5.71%, respectively) (Fig. 9a). The genes related to nitrogen (13.67%) and sulfur (16.47%) metabolism remained predominantly abundant in CM-microbiomes (Fig. 9a). Furthermore, within OP metabolic pathways, the predicted gene families of NADH dehydrogenase (*ndh*), ATPase subunits (*atp*A and *atp*D), and inorganic pyrophosphatase (*ppa*) had higher relative abundances in CM-microbiomes. The gene and/or protein families associated with bacterial chemotaxis (*Che*C*, Che*D*, Che*R*, Che*X*, Che*Z*, mot*A*, mot*B) were predominantly enriched in SCM- pathogens (57.89%), and those encoding for methyl-accepting chemotaxis (*tsr, tar, trg* and *tap* genes) had higher relative abundances in RCM (5.56%) and CM (4.13%)-causing pathogens. Notably, genes responsible for bacterial flagellar assembly (41.30%), succinyl-CoA synthetase alpha-beta subunits (*suc*C, *suc*D) (31.25%), and two-component systems (*Che*A*, Che*B*, Che*BR*, Che*V*, Che*Y) (18.52%) remained overexpressed in RCM-microbiomes, and genes coding for p53 signaling pathway had relatively higher expression in SCM (1.56%)- causing microbiomes (Fig. 9a). Conversely, the H-milk microbiomes had comparative overexpression in genes coding for pyruvate metabolism (19.31%), cytochrome c oxidase subunits such as *cox*A, *cox*B and *cox*C (5.47%), and protoheme IX farnesyltransferase, *cyo*E (4.77%) (Fig. 9a, Supplementary Data 2).

We also sought to gain further insight into the SEED hierarchical protein functions represented by different genes, and found 25 statistically different (p = 0.001, Kruskal-Wallis test) SEED functions in mastitis (CM, RCM, SCM) and H-milk metagenomes. Overall, the mastitis associated microbiome showed a higher relative abundance of these SEED functions compared to H-milk microbiomes, except for plasmid related functions (highest in H microbes; 6.90%) (Fig. 9b, Supplementary Data 2). The BarA-UvrY (*Sir*A)-two-component regulatory system was found as the most abundant functional pathway in RCM-pathogens (100.00%) followed by CM (85.60%), H (83.66%) and SCM (33.33%)-milk microbiomes (Fig. 9b, Supplementary Data 2). The CM-microbiomes were enriched in genes coding for phage integration and excision (36.16%), autoinducer-2 (AI-2) transport and processing (*lsr*_ACDBFGE_ operon, 34.74%), pathogenicity islands (15.28%), *Yjg*K cluster-linked to biofilm-formation (15.04%), NADPH quinone oxidoreductase-2 (3.34%) and *Tyr*R associated virulence (2.93%) compared to the microbes of the other metagenomes. The RCM-microbiomes however had a higher abundance of SEED functions involved in biofilm adhesin biosynthesis (80.0%), quorum-sensing and biofilm-formation (29.41%), glutathione non-redox reactions (20.83%), regulation of oxidative stress (ROS) response (12.50%), transposable elements (12.06%), phage DNA synthesis (2.38%), and gene transferring agent (1.89%).The SCM-microbiomes particularly showed the elevated expression of genes related to ROS (33.33%), oxidative stress (28.57%), virulence regulation (19.35%), phage related functions such as *r1t*-like *Streptococcal* phages (35.59%), phage regulated gene expression (12.71%), phage packaging machinery (11.86%), phage replication (9.32%) and prophage lysogenic conversion (2.54%) modules. In contrast, most of these SEED modules had relatively lower abundance in H-milk microbiomes (Fig. 9b, Supplementary Data 2).

## Discussion

The current study represents the first ever proof-of-concept to decipher dynamics changes in microbiome composition, and abundances in CM, RCM, SCM and H-milk samples through the cutting-edge WMS technology. The present findings of higher taxonomic resolution, and predicted protein functions, partially consistent with our previous report^6^, and many of the recent 16S rRNA partial gene-based studies^13, 14, 25^. The microbiome diversity (alpha and beta diversity) measures provided that microbial dysbiosis is closely linked to different stages of mastitis. Compared to our previous study^6^, we observed increased microbial diversity and species richness in CM, RCM, and SCM-metagenomes than the healthy-controls. Beta diversity^6, 20^ also revealed a substantial microbial disparity between healthy-controls, and mastitis-milk samples keeping the closest relationship of the CM-microbiomes to RCM followed by SCM, and H-microbiomes. Regardless of higher taxonomic abundances, the bovine mastitis-associated microbiome remained inconsistent, and fluctuates more within CM, RCM, and SCM-metagenomes than those of H-milk metagenome, agreeing with several recent studies^6, 11, 12^.

In bovine mastitis, predominantly identified pathogens are bacteria^6, 7, 10–16^, however, other concomitant microbial players like archaea and viruses^6, 12^ could also be detected, highlighting the novel insights from metagenomics compared to amplicon sequencing. The bovine milk microbiome was dominated by *Proteobacteria*, *Firmicutes*, *Actinobacteria* and *Bacteroidetes*, and their relative abundance also varied across the samples of four metagenomes. These results are in line with bovine^6, 7, 11, 15^ and human^20^ mastitis microbiome reported earlier. Previously, *Proteobacteria* and *Firmicutes* were reported as the major phyla in bovine SCM^15^ supporting our current findings. During the pathophysiology of mastitis, mammary epithelial cells reduce their capacity to beta-oxidation which alters the composition of indigenous microbiota, reducing the number of anaerobic *Firmicutes*, and allowing the proliferation of different genera and/or strains belonged to *Proteobacteria* blooms^7, 11, 12, 33, 34^.

Metagenomics^6, 7, 13–15^ and isolate sequencing^35^ of mastitis pathogens provided overwhelming evidence that microbial traits, for instance, pathogenicity, virulence, antibiotic resistance, and metabolic potentials are linked with strain-specific genomic characteristics^21, 36–38^. In this study, different strains of *Acinetobacter* were detected as the predominantly abundant CM and RCM-causing pathogen, as also previously reported in human^39^, bovine^6^ and bubaline^40^ mastitis. The CM and RCM-metagenomes also had a higher abundance of strains of the *Pseudomonas* which can potentially cause opportunistic infections^6, 12^. However, in SCM- metagenome, the abundance of different strains of *Acinetobacter* and *Pseudomonas* remained substantially lower compared to CM and RCM-metagenomes. We also revealed that different strains of *Escherichia*, *Aeromonas*, *Klebsiella*, *Shigella*, *Enterobacter*, *Salmonella*, *Pantoea*, *Citrobacter*, *Shewanella*, *Lactococcus*, and *Cronobacter* routinely colonize the mammary tissue or quarters, and manifest different classes of mastitis^6, 12, 24^. The SCM-metagenome was predominated by different strains of *Lactococcus*, *Chryseobacterium*, *Ralstonia*, *Serratia*, *Riemerella*, *Streptococcus*, and *Pedobacter* corroborating the findings of several earlier reports^11, 12, 15^. Many of these microbiotas may act as potential opportunists by interfering with metabolism, host defense, and immune development^41–44^ to producing different magnitude of udder infections. Our present findings could be further supported by the potential existence of endogenous entero-mammary pathway, and through this axis the gut or rumen microbiomes migrate to the mammary gland to manifests different episodes of mastitis^6, 24, 38^. The healthy-milk metagenome was also dominated by different strains of environmental, gut and animal skin originating microbes. Though the pathogenic mechanisms of these commensal microbes are largely unknown, they can cause opportunistic infections in the mammary glands and/or quarters with or without varying degrees of clinical episodes by producing different virulence factors^45^ particularly in immunocompromised hosts^6, 24, 42, 43^.

The state-of-the-art WMS approach also provided an exciting opportunity for investigating the integrated cross-kingdom interactions of ‘multibiome’– traditionally neglected components of the milk microbiome such as archaea and viruses (< 1.0% genes coding for total microbiome)^46, 47^. Unlike bacteria, the diversity and composition of archaea^46^ and viruses^47^ always remained much lower in both mastitis and healthy-milk^6^. The archaeal fraction of the microbiome is mostly represented by the methanogenic and thermophilic genera of *Euryarchaeota* in CM and H-milk, and halophilic archaeal genera in SCM and RCM- metagenomes. Most of the viruses detected in both mastitis and healthy-milk metagenomes are tailed bacteriophages capable of infecting bacteria, and archaea during physiological challenges that can compromise the defense mechanisms of the udder particularly in lactating cows^6, 7^. Even though the role of these accompanying microbiotas in the pathophysiology of bovine mastitis has not been understood within the frames of a typical host-pathogen interaction, these microbiotas could cease the opportunity during the pathological changes in the mammary glands created by bacteria^46, 47^. They may occupy the microenvironments suitable for anaerobic metabolism, make way for opportunistic pathogens (pathobionts), already present in milk, and/or gain secondary access to the mammary gland when the host is in unnatural condition or immune compromised^20, 21^, and follow the similar virulence mechanisms of bacterial pathogens producing more severe and prolonged recurrent mastitis (RCM)^46, 47^. Furthermore, a higher abundance of *Haloquadratum* and *Natronomonas* in SCM and RCM-milk may possess some accessory capsular switching genes for their adaptation to environmental fluctuations like bacterial pathogens to causing different magnitudes of diseases^48^.

The mastitis-associated pathogens harbored a wide range of VFGs, and bacteria having higher abundances also possessed an increased number of associated VFGs in the corresponding metagenomes. The CM, RCM, and SCM-microbiomes had a higher number of VFGs related to innate-immune responses, pyoverdine or antibiotics susceptibility altering, twitching motility and chemotaxis, adherence to mammary epithelial cells and invasion, two-component system, flagellar assembly, intracellular survival and multiplication, bacteriocins production and cytotoxic activity, energy transduction, capsular proteins, lipid biosynthesis, ABC transporter, biofilm-formation (BF), and quorum-sensing, and siderophores linked to pathogenesis^1, 8, 24, 50–53^. Conversely, H-milk microbiomes harbored a higher number of genes required for innate-immune responses, intracellular survival in competition, pili and flagella expression, bacterial chemotaxis, alginate biosynthesis, and BF which play important roles in protecting bacteria from the host-defenses^8, 50–53^. The underlying mechanisms for microbial colonization in mammary tissue modulated by these VFGs are not well established. However, VFGs can enable microbiome to overcome host defense^8^, immune-mediated colonization of the mammary tissues by suppressing the regrowth of resident commensal microbiota^7, 24, 49^, and subsequent inclusion of opportunistic pathogens with the progression, and recurrence of the disease^7, 24, 49, 54, 55^.

We found various homologs of ARGs belonging to different protein families among the microbiome of four metagenomes, and the composition and abundances of the ARGs remained significantly correlated with the abundances of the related bacteria. Furthermore, the ARGs also varied greatly in different states of mastitis (CM, RCM, SCM) supporting the dynamic dysbiosis of microbiomes in the corresponding metagenomes, their genetic diversity, and selective pressure for the maintenance of ARGs^9, 55^. The β -Lactamases, major resistance determinant for β -lactam antibiotics in Gram-negative bacteria^56^, remained with extended-spectrum in mastitis microbiome. However, β -lactamase resistant gene of OXA (oxacillinase) group, *bla*_OXA_ remained predominantly abundant in H-milk microbes. This ARG which might be involved in hydrolytic activity against multiple antibiotics^9^, and sometimes augmented in mastitis pathogens by additional resistance mechanisms of impermeability or efflux system^57^. The tetracyclines resistant genes were found only in CM, RCM and SCM-pathogens, with comparatively higher abundances in SCM-microbiome, and corroborated with several earlier reports^56, 57^. In addition, aminoglycosides, trimethoprim, florfenicol, and sulfonamide-resistant genes were predominantly abundant in CM-microbiome, which could be due to drug-specific acquired resistance mechanisms^56^. Higher number and classes of ARGs including macrolides, fosfomycin, and quinolones resistance genes in RCM-microbiome might be resulted from their infrequent use, and subsequent treatment failure in bovine CM. The possible mechanisms for the detected ARGs include enzymatic inactivation, prevention of antibiotic-bindings to the targets, catalytic activity, folate pathway antagonist-attributed resistances, and efflux pump and/or system conferring resistance to a wide range of antibiotics^9, 56, 58^. These ARGs can easily travel and spread throughout different dairy environments because of wind and runoff waters, climate change, human activities and contact with wild animals, in particular, migratory ones^9, 56^.

The present study found several important predicted metabolic functions that altered in their abundances between CM, RCM, SCM and healthy-milk microbiome as also reported previously in lactating cows^13^, women^20^ and mouse^44^. Our analysis revealed that similar metabolic features of microbiota between different states of mastitis are associated with their early colonization, and disease persistent and/or progression^7, 17, 18^. Recent studies have shown that metabolites are involved in modulating immune function^37, 51^. The mastitis-associated microbiomes harbored a higher number of genes for nutrient transport^60^, and carbohydrate metabolism through TCA-cycle^61^. Moreover, increased benzoate degradation by different strains of *Acinetobacter* and *Klebsiella* through TCA-cycle might promote bacterial growth, and virulence factors expression during mammary gland pathogenesis^62^. The methyl-accepting transmembrane chemotaxis genes were upregulated in RCM and CM-associated pathogens suggesting their role in cell-to-cell communication, early phase attachment to or entry into the udder tissues, and virulence regulation^62^. The aerotaxis receptor gene was overexpressed in H milk microbiomes, which is thought to sense internal redox states, and mediate bacterial movement towards optimal oxygen conditions or mediate maximal growth of bacteria^24, 62^. The p53 signaling being remained increasingly identified in healthy-milk microbiomes might response to a stress signal, and activated in a specific manner by post-translational modifications which leads to either cell cycle arrest or mammary gland apoptosis^63^. The global regulating unit BarA-UvrY (*Sir*A)-two-component system is well conserved in many bacterial species (γ- *Proteobacteria* in particular), and regulate the production of extracellular factors (exoenzymes or toxins), quorum-sensing, motility, and diverse metabolic functions^51, 64^. In many mastitis-causing microbiomes, BarA/UvrY is important for regulation and coordination of pathogenicity and group behaviors^51, 64^. The relative overrepresentation of filamentous phage-related genes in mastitis-pathogens may suggest that bacteriophages participate in the horizontal gene transfer among the members of bovine milk microbiome and eventually to mammary gland pathogens^65^. The altered metabolic pathways such as transport and processing, pathogenicity islands, and BF in mastitis-pathogens are associated with a diverse array of virulence mechanisms for mammary gland pathogenesis^24^. Bacterial BF is a strain-specific or genetically-linked trait with selective advantage in pathogenesis of mastitis and harmful to mammary tissues, since they can promote the phagocyte release, proliferation of reactive oxygen and nitrogen species, and transfer of antibiotic resistance^6, 9^. Moreover, RCM-microbiomes had overexpression of genes encoding biofilms adhesins^24, 50^, glutathione non-redox reactions^66^, regulation of oxidative stress^67^, and transposable elements^68^ which play an important role in many chronic-recurrent bacterial infections like bovine RCM.

## Conclusions

The shotgun metagenome sequencing of milk samples from bovine mastitis (CM, RCM, SCM) and healthy quarters revealed a complex microbial diversity, with dynamic alteration of microbial communities, and its association with the functional determinants in different pathophysiological categories of bovine mastitis. We demonstrated that microbial dysbiosis in different types of mastitis resulted in the depletion of beneficial microbes and enrichment of the opportunistic pathogens. Our results suggest that co-occurrence of VFGs and ARGs possibly playing determining roles in the adaptation of different microbiomes mastitis and healthy milk. Several imputed functional pathways differed between mastitis and healthy controls, perhaps reflecting metabolic changes associated with the progression of mastitis pathogenesis. These findings point toward the development microbiome-based diagnostics and therapeutics for this economically burdened disease. However, future studies should delve further into characterizing microbiome diversity with larger sample sizes and the inclusion of gut/rumen microbiome sampling in addition to the milk samples for direct testing of microbial transfer across this axis to confirm the shift of microbiome and associated functions.

## Methods

### Screening for mastitis, and sampling

Details of the study population and collected samples are presented in Supplementary Table 1. The California Mastitis Test (CMT^®^, Original Schalm reagent, ThechniVet, USA) was used to screen different states of mastitis (clinical mastitis, CM; recurrent clinical mastitis, RCM; subclinical mastitis, SCM)^19^. Cows with CM (mild, moderate, and severe) typically have abnormalities in the milk such as clots and flakes, swelled, red and hard mammary gland or systemic illness. The RCM can be either a persistent CM of the bovine mammary gland by a mastitis pathogen or a reinfection of a quarter or udder after bacteriological cure of CM while the SCM is inflammation of the mammary gland that does not create visible changes in the milk or the udder^6, 11, 14–19^.

A total of 20 milk samples (CM = 5, RCM = 6, SCM = 4, and healthy (H) = 5) from 20 lactating cows at early stages of lactation (within 10 - 40 days of calving) were collected from five districts in Bangladesh: Gazipur = 8 (24.19 N, 90.47 E), Dhaka= 3 (23.81 N, 90.41 E), Manikgonj = 3 (23.86° N, 90.00° E), Chattogram = 4 (22.20 N, 91.98 E), Sirajgonj = 2 (24.31° N, 89.57° E). The sampling patterns followed the collection of one mastitis and one H milk samples from the same farm. Approximately 15-20 ml of milk from each cow was collected in a sterile falcon tube during morning milking (8.0-10.0 am), with an emphasis on pre-sampling disinfection of teat-ends and hygiene during sampling^14, 19^. CM cows which did not recover fully in spite of receiving different conventional antibiotic treatments, with subsequent persistence of CM episodes, are defined as the cows with RMC^18^. The protocol for milk sample collection was approved by the Animal Experimentation Ethical Review Committee (AEERC) of the University of Dhaka under reference number 79/Biol.Scs. The milk samples were then transported to the laboratory and stored at -20°C until DNA extraction.

### DNA extraction, shotgun metagenomic sequencing, and sequence reads preprocessing

The total genomic DNA (gDNA) content of each milk sample was extracted by an automated DNA extraction platform (Promega, UK) using previously published protocols^6^ along with the manufacturer’s instructions. The quantity and purity of the extracted DNA was measured with NanoDrop (ThermoFisher, USA) by measuring 260/280 absorbance ratios. Shotgun whole metagenome sequencing (WMS) libraries were prepared with Nextera XT DNA Library Preparation Kit^69^ and paired-end (2×150 bp) sequencing was performed on a NextSeq 500 machine (Illumina Inc., USA) at the George Washington University Genomics Core facility. The current metagenomic DNA generated 416.64 million reads with an average of 20.83 million (maximum = 39.75 million, minimum = 2.72 million) reads per sample (Supplementary Data 1). The resulting FASTQ files were concatenated and filtered through BBDuk^6^ (with options k=21, mink=6, ktrim=r, ftm=5, qtrim=rl, trimq=20, minlen=30, overwrite=true) to remove Illumina adapters, known Illumina artifacts and phiX. Any sequence below these thresholds or reads containing more than one ‘N’ were discarded. On average, 19.28 million reads per sample (maximum = 37.36 million, minimum = 2.50 million) passed the quality control step (Supplementary Data 1).

### Taxonomic mapping, classification, diversity, and community analysis

Shotgun whole metagenome sequencing (WMS) data were analyzed using both mapping-based and assembly-based hybrid methods of PathoScope 2.0 (PS)^28^ and MG-RAST (MR)^29^, respectively. In PS analysis, a ‘target’ genome library was constructed containing all bacterial sequences from the NCBI Database (https://en.wikipedia.org/wiki/National_Center_for_Biotechnology_Information) using the PathoLib module^28, 69^. The reads were then aligned against the target libraries using the Bowtie 2 algorithm^70^ and filtered to remove the reads aligned with the cattle genome (bosTau8) and human genome (hg38) as implemented in PathoMap (very sensitive local -k 100 --score-min L,20,1.0). Finally, the PathoID^71^ module was applied to obtain accurate read counts for downstream analysis. In these samples, 16.88 million reads (4.05% of total reads) were mapped to the target reference genome libraries after filtering the cow and human genome (Supplementary Data 1). The raw sequences were simultaneously uploaded to the MR server (release 4.0) with properly embedded metadata and were subjected to optional quality filtering with dereplication and host DNA removal, and finally screening for taxonomic and functional assignment. Alpha diversity (diversity within samples) was estimated using the observed species, Chao1, ACE, Shannon, Simpson, and Fisher diversity indices^72^ for both PS and MR read assignments and counts. To visualize differences in bacterial diversity, principal coordinate analysis (PCoA) based on the Bray-Curtis distance method^73^ for MR data (genus level), and non-metric multidimensional scaling (NMDS) measured by weighted-UniFrac distance on PS data (at strain level) through Phyloseq R package (version 3.5.1)^74^ were performed. Taxonomic abundance was determined by applying the ‘‘Best Hit Classification’’ option using the NCBI database as a reference with the following settings: maximum e-value of 1×10^-30^; minimum identity of 95% for bacteria, 60% for archaea and viruses, and a minimum alignment length of 20 as the set parameters. The phylogenetic origin of the metagenomic sequences was projected against the NCBI taxonomic tree and determined by the lowest common ancestor (LCA) with the same cutoff mentioned above.

### Virulence factors and antibiotic resistance-associated genes

The virulence factor database (VFDB) and DNA sequences with full datasets for virulence factors of pathogenic bacteria^30^ were used to identify virulence factors associated genes (VFGs) among the microbiomes of the four metagenomes. Similarly, to identify the antibiotics resistance genes (ARGs) among the microbiomes of all metagenomes, we used the ResFinder 2.0 database^31^. Both VFDB and ResFinder databases have been integrated within AMR++ pipeline^31^ to identify the respective genes and/or protein families. Every protein included in each of the addressed metagenomes was used as a query to search for similarities to either VFGs or ARGs protein-coding traits. Hence, we aimed at retrieving the best hit (best-scored alignment) that enabled us to assign a VFG or ARG function to each of the aforementioned metagenomic proteins. Furthermore, we computed the representative counts for the different gene families that coded for either VFGs or ARGs traits from all the generated alignments between our metagenomic query cohort and the preceding databases. Thus, the amounts of different classes (gene families) that are present in each metagenome represent its diversity in terms of VFGs or ARGs traits. We used OmicCircos (version 3.9)^75^, an R package based on a Python script for circular visualization of both diversity and composition of virulence factors associated genes (VFGs) across the four metagenomes under study.

### Functional profiling of the microbiomes

To perform taxonomic functional classification, the reads were mapped onto the Kyoto Encyclopedia of Genes and Genomes (KEGG) database^32^, and SEED subsystem identifiers, respectively, on the MR (version 4) metagenome analysis server^29^ using the partially modified set parameters (e*-*value cutoff: 1×10^-30^, min. % identity cutoff: 60%, and min. alignment length cutoff: 20).

### Statistical analysis

The non-parametric test Kruskal-Wallis rank sum test was used to evaluate differences in the relative percent abundance of taxa in mastitis (CM, RCM, SCM) and H groups for PS data. Comparative taxonomic and functional profiling was performed with the reference prokaryotic metagenomes available in MG-RAST database for statistical analyses. The gene counts were normalized by dividing the number of gene hits to individual taxa/function by total number of gene hits in each metagenome dataset to remove bias due to differences in sequencing efforts. To identify differentially abundant SEED or KEGG functions, virulence factors (VFGs), and antimicrobial resistance (ARGs) in the metagenomes, statistical tests were applied with non-parametric test Kruskal-Wallis rank sum test at different KEGG and SEED subsystems levels IBM SPSS (SPSS, Version 23.0, IBM Corp., NY USA).

### Data availability

The sequence data reported in this paper are available in the NCBI database (BioProject PRJNA529353) and European Nucleotide Archive (BioProject, PRJEB31210).

### Ethics Statement

The protocol for milk sample collection from lactating dairy cows was approved by the Animal Experimentation Ethical Review Committee (AEERC), Faculty of Biological Sciences, University of Dhaka (under reference number: 79/Biol.Scs).

## Supporting information

Main Figure Legends

## Author contributions

MNH, MS, and MAH conceived and designed the study. MNH surveyed and collected field samples. MNH, SW and AI carried out laboratory works including DNA extractions and sequencing. MNH, AI, MSR, MRI and KAC conceived, designed, and executed the bioinformatics analysis. MNH interpreted the results, visualized the figures, and drafted the manuscript. MAH and KAC contributed chemicals and reagents. AA, MS, AMAMZS, KAC and MAH contributed intellectually to the interpretation and presentation of the results. Finally, all authors have approved the manuscript for submission.

## Acknowledgements

The Bangladesh Bureau of Educational Information and Statistics (BANBEIS), Ministry of Education, Government of Bangladesh (GoB) (Grant No. LS2017313) supported this work. The first author of this manuscript, MNH received his PhD Fellowship from the Bangabandhu Science and Technology Fellowship Trust, Ministry of Science and Technology, GoB. The authors thank Rebecca A. Clement and Stephanie Warnken, PhD students at the Computational Biology Institute, Milken Institute School of Public Health, The George Washington University, USA for their technical support in sequencing and bioinformatics. We also acknowledge the cooperation of the dairy farmers for allowing us to conduct the study in their farms.

## Additional Information

Supplementary information supporting the findings of the study are available in this article as Supplementary Data files, or from the corresponding author on request.

---

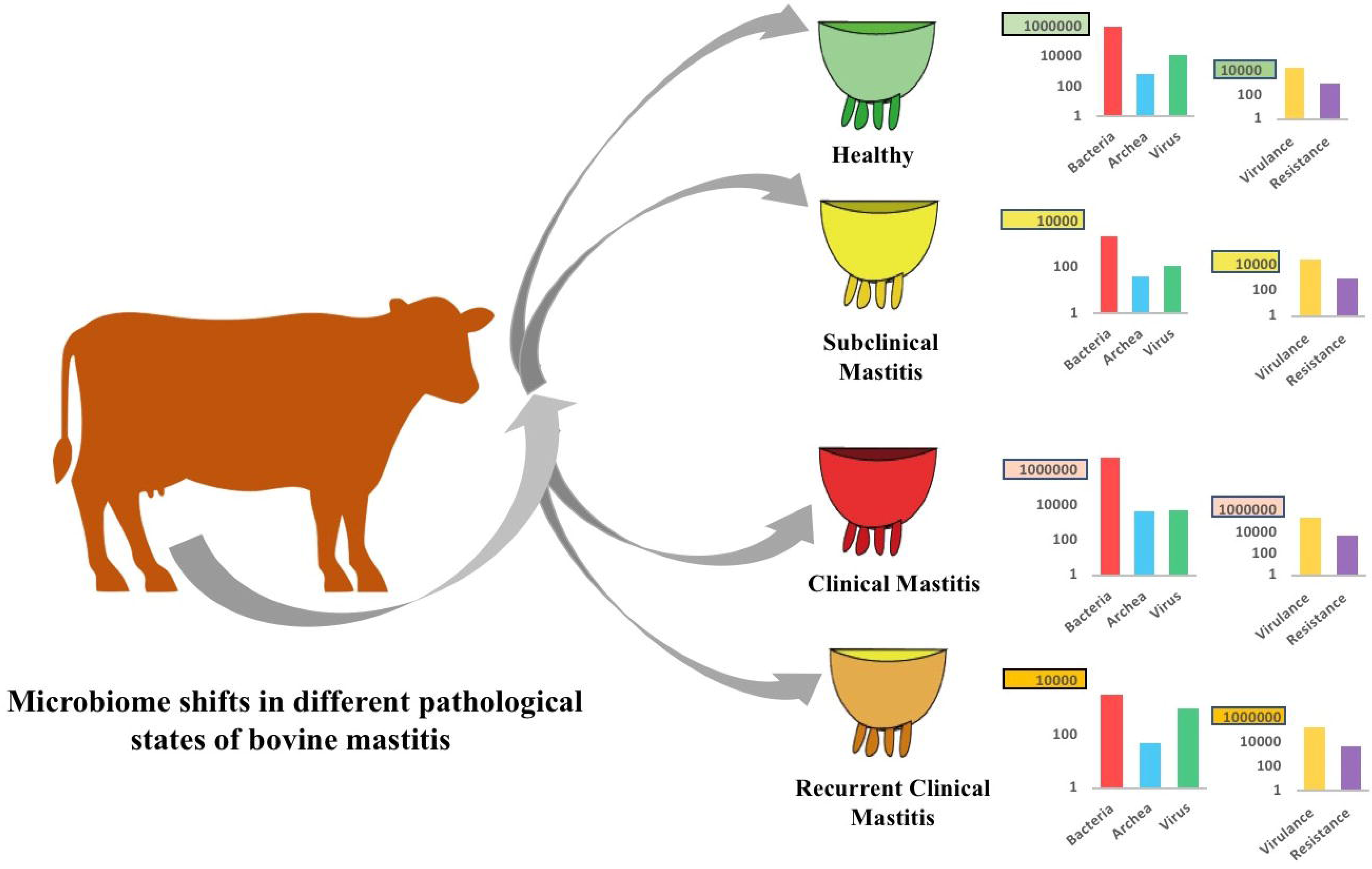

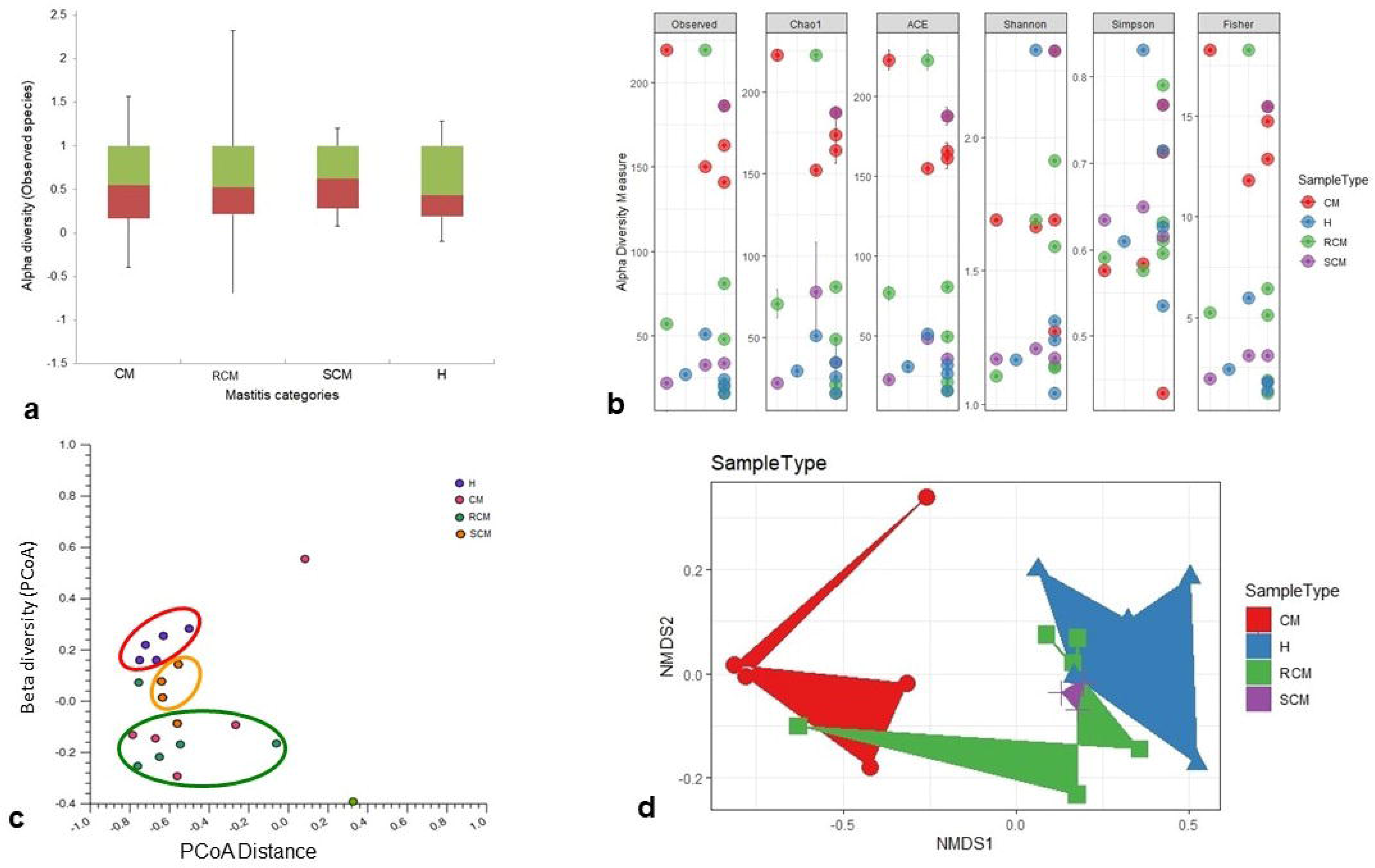

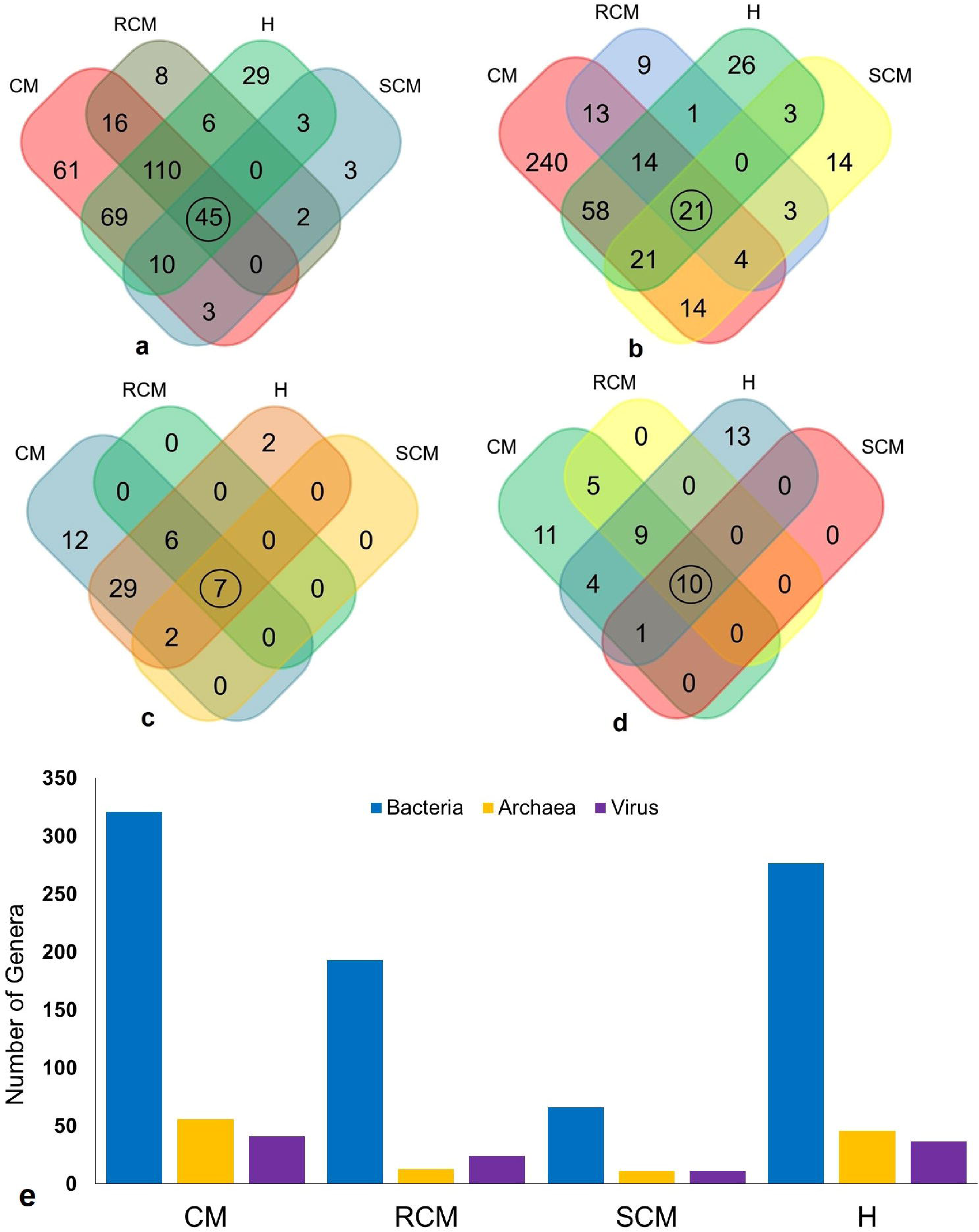

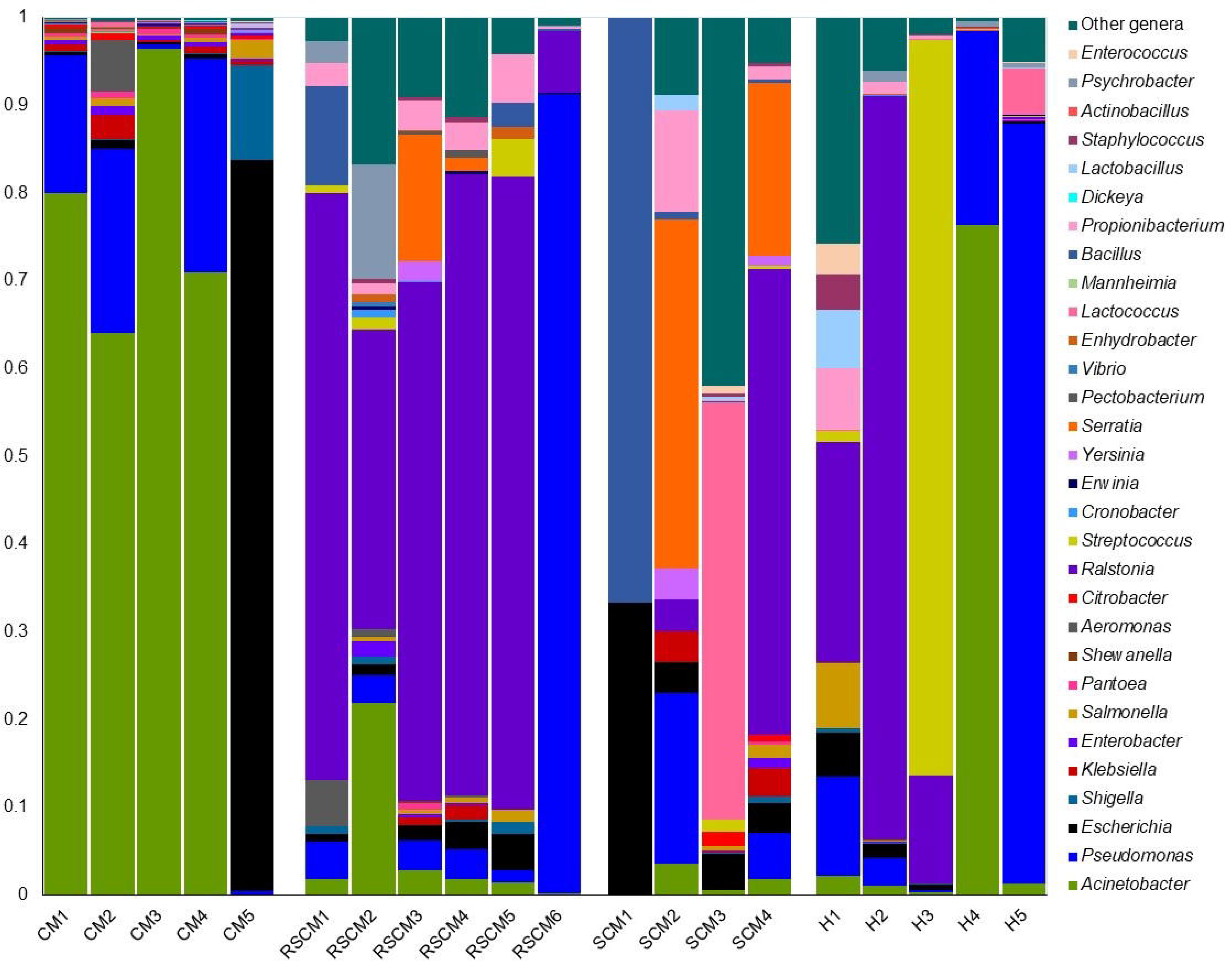

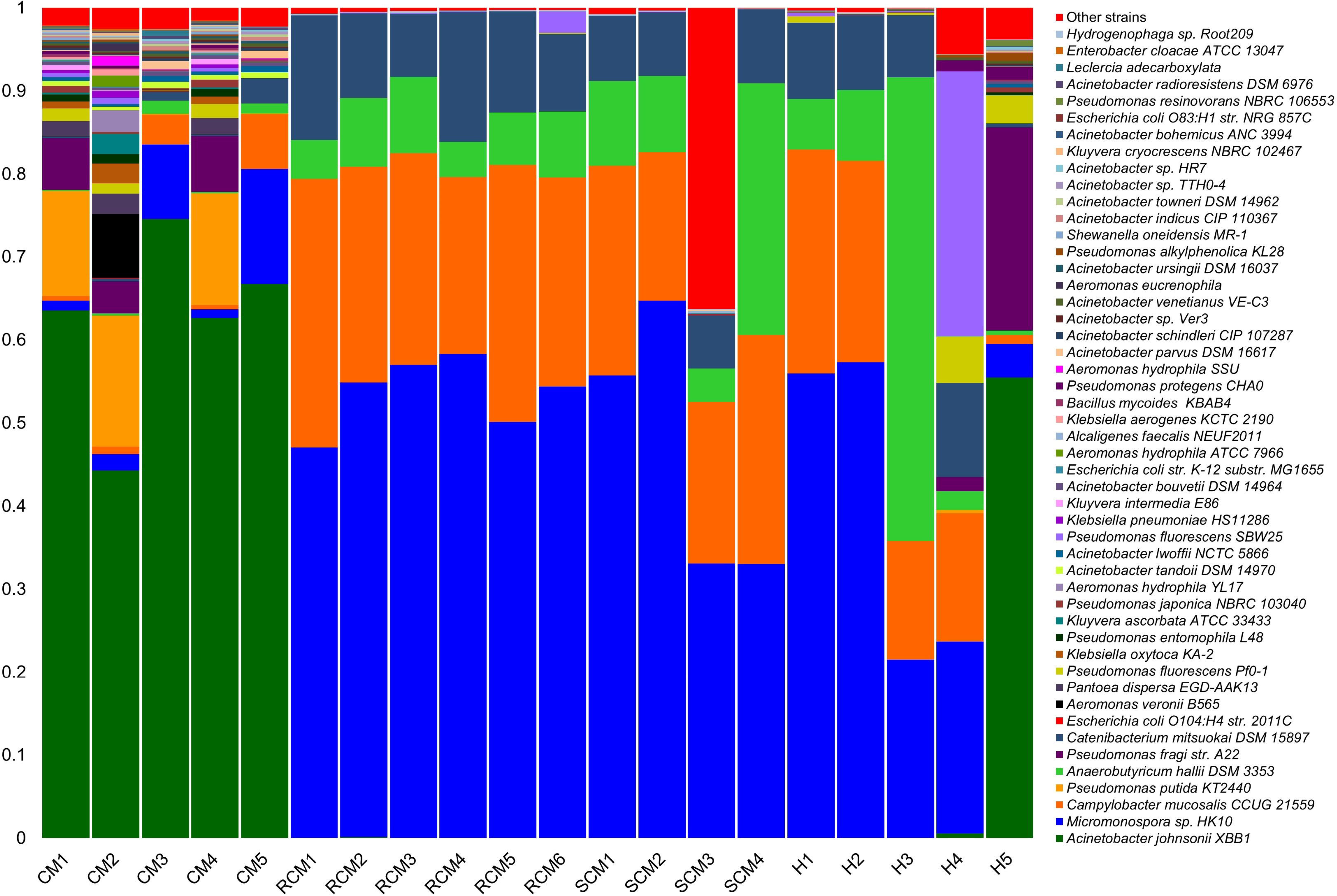

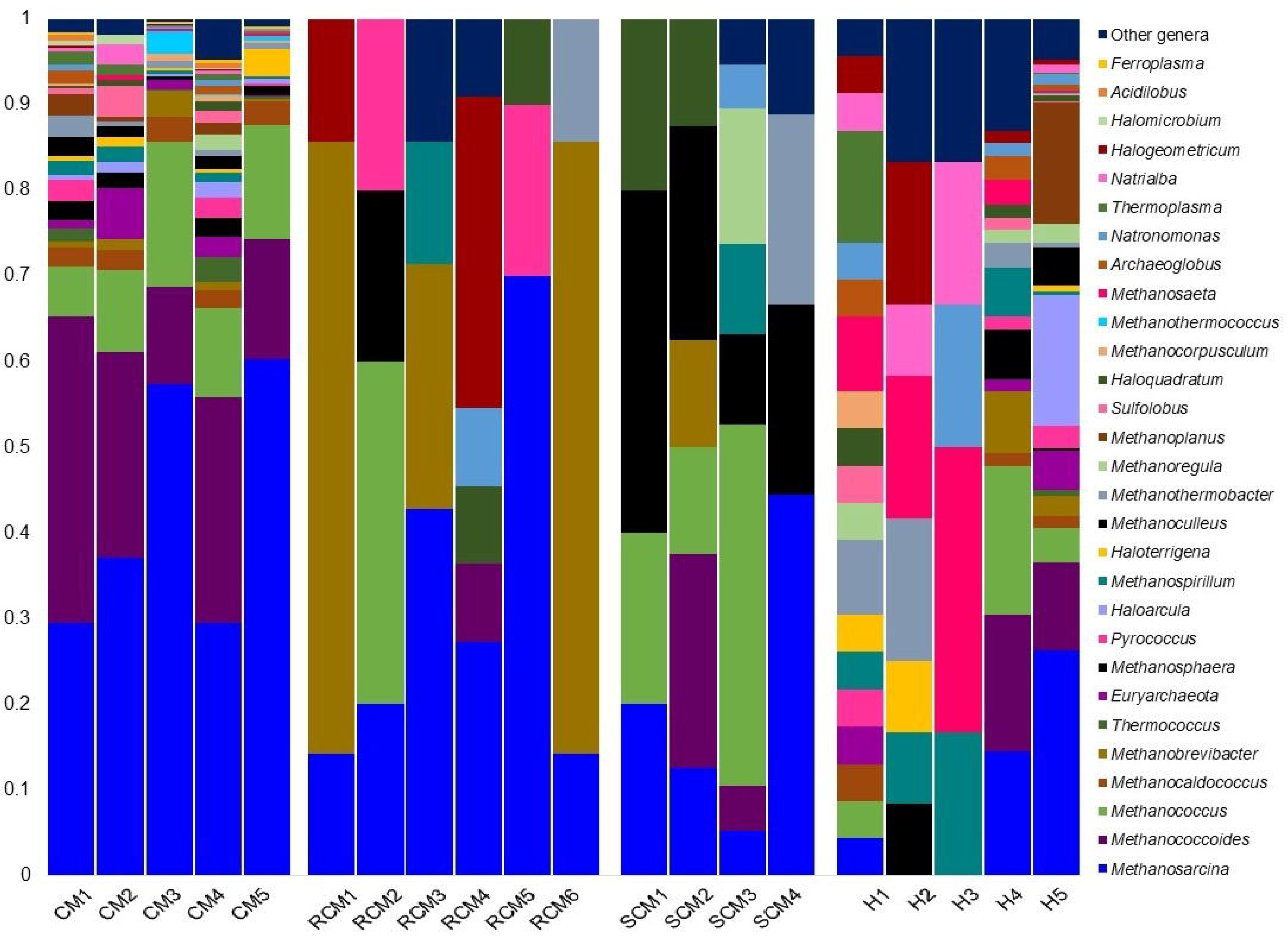

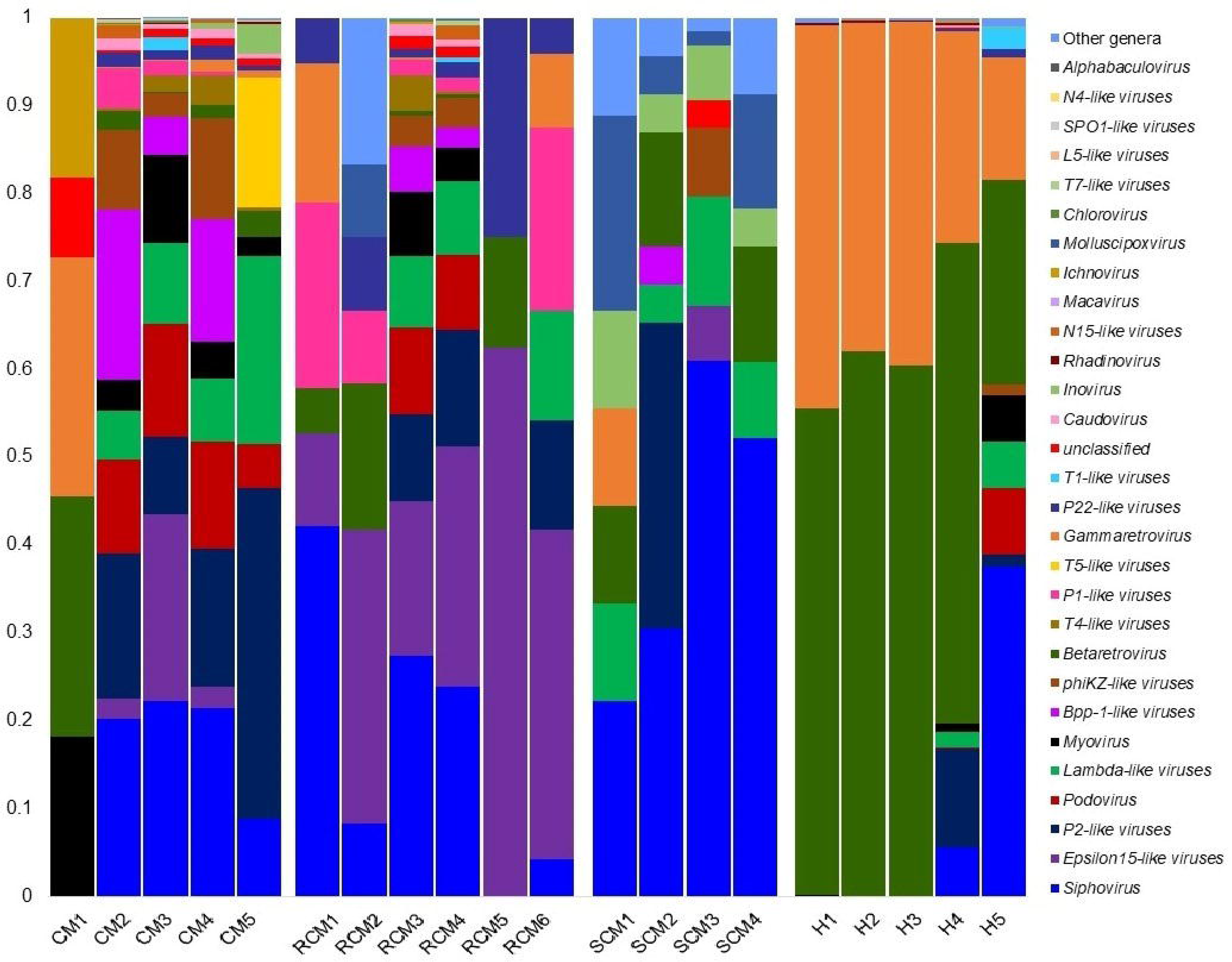

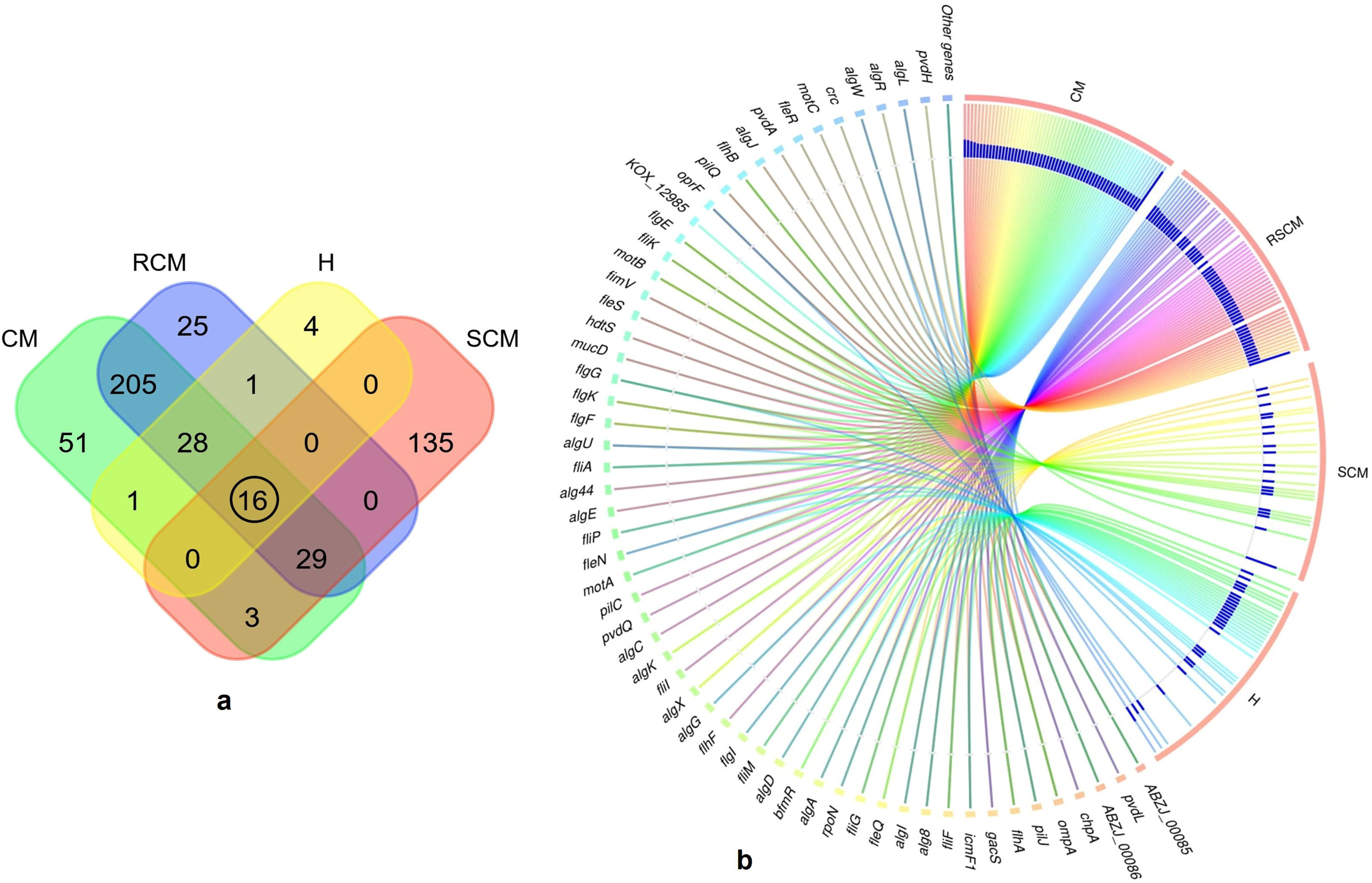

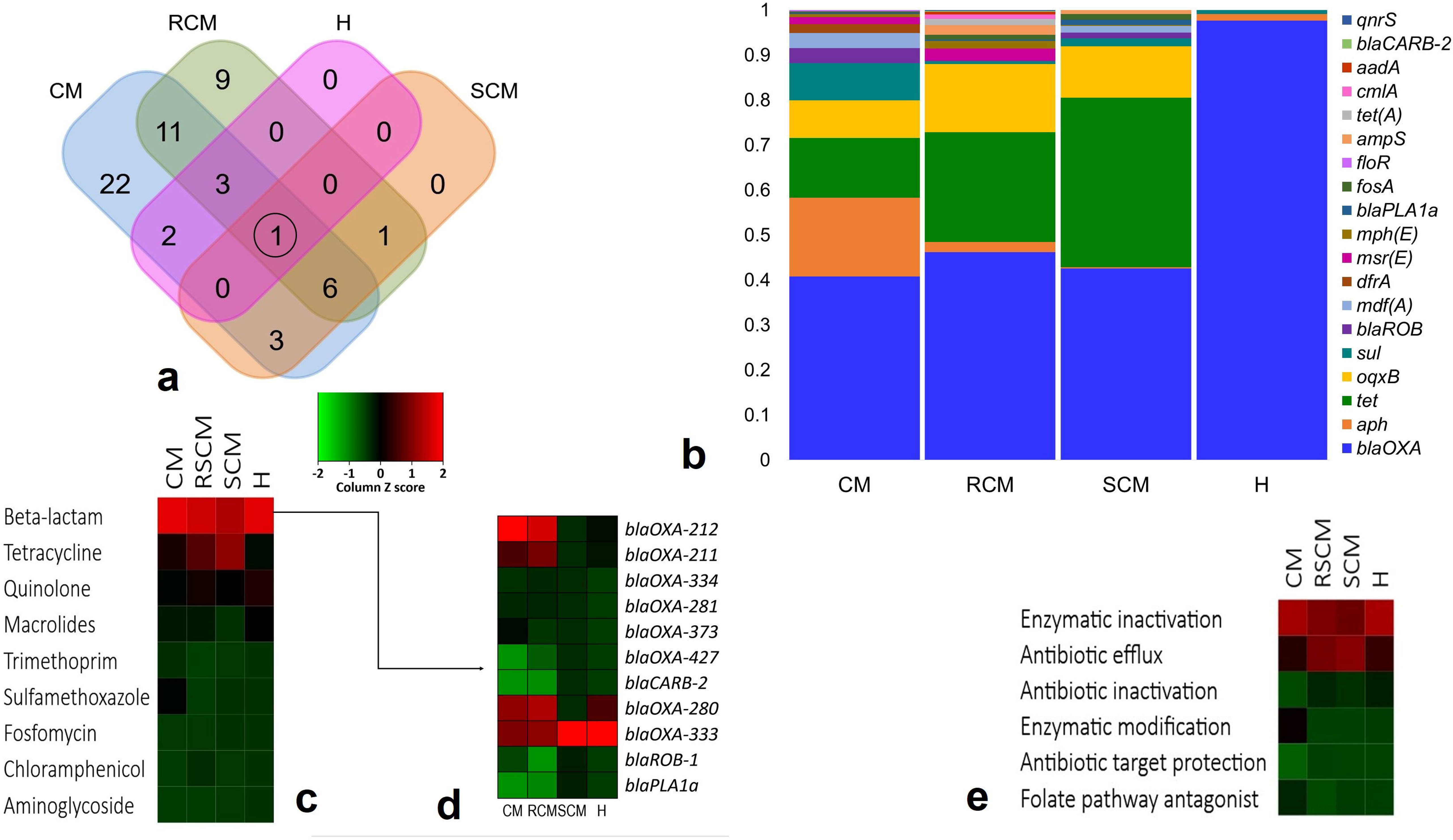

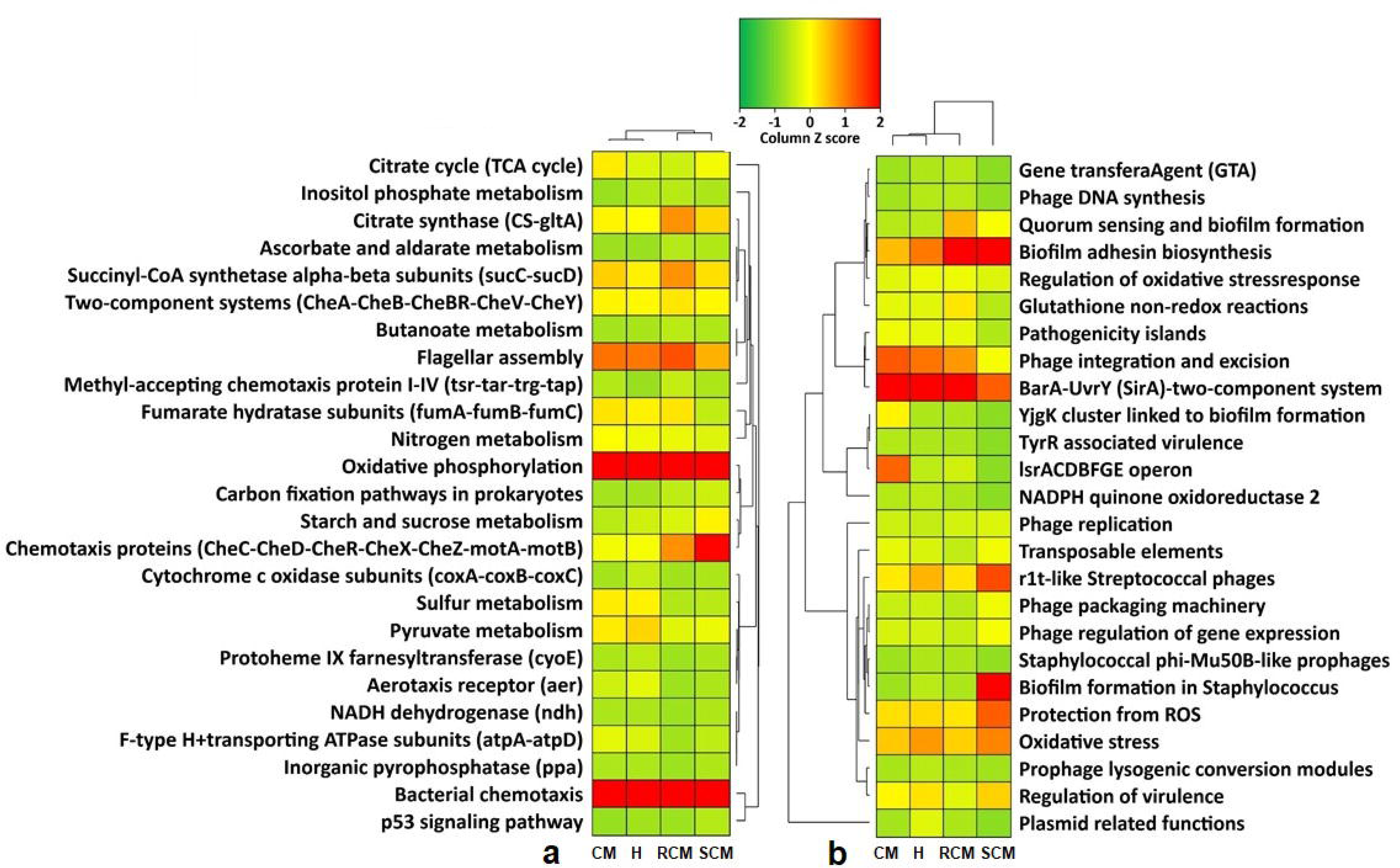

## References

1. Thompson-Crispi, K., Atalla, H., Miglior, F. & Mallard, B. A. Bovine mastitis: Frontiers in immunogenetics. Front. Immunol. 5, 493 (2014).

2. Reyes-Jara, A., Cordero, N., Aguirre, J., Troncoso, M. & Figueroa, G. Antibacterial Effect of Copper on Microorganisms Isolated from Bovine Mastitis. Front. Microbiol. 7, 626 (2016).

3. Jiménez, E. et al. Metagenomic analysis of milk of healthy and mastitis-suffering women. J. Human Lactation 31, 406–415 (2015).

4. Mediano, P. et al. Microbial diversity in milk of women with mastitis: potential role of coagulase-negative *staphylococci*, viridans group *streptococci*, and *corynebacteria*. J. Human Lactation 33(2), 309–318 (2017).

5. Sakwinska, O. & Bosco, N. Host microbe interactions in the lactating mammary gland. Front. Microbiol. 10, 1863 (2019).

6. Hoque, M. N. et al. Metagenomic deep sequencing reveals association of microbiome signature with functional biases in bovine mastitis. Sci. Rep. 9, 13536 (2019).

7. Derakhshani, H. et al. Invited review: microbiota of the bovine udder: contributing factors and potential implications for udder health and mastitis susceptibility. J. Dairy Sci. 101(12), 10605–10625 (2018).

8. Thomason, C.A. et al. Resident Microbiome Disruption with Antibiotics Enhances Virulence of a Colonizing Pathogen. Sci. Rep. 7, 16177 (2017).

9. Escudeiro, P., Pothier, J., Dionisio, F. & Nogueira, T. Antibiotic Resistance Gene Diversity and Virulence Gene Diversity Are Correlated in Human Gut and Environmental Microbiomes. mSphere 4(3), e00135–19 ((2019).

10. Hoque, M. N. et al. Insights into the Resistome of Bovine Clinical Mastitis Microbiome, a Key Factor in Disease Complication. Front. Microbiol. 11, 860 (2020). doi: 10.3389/fmicb.2020.00860.

11. Oikonomou, G. et al. Microbiota of cow’s milk; distinguishing healthy, sub-clinically and clinically diseased quarters. PLoS One 9(1), e85904 (2014).

12. Oikonomou, G. et al. Milk microbiota: what are we exactly talking about?. Front. Microbiol. 11, 60 (2020).

13. Cremonesi, P. et al. Milk microbiome diversity and bacterial group prevalence in a comparison between healthy Holstein Friesian and Rendena cows. PLoS One 13(10), e0205054 (2018).

14. Falentin, H. et al. Bovine teat microbiome analysis revealed reduced alpha diversity and significant changes in taxonomic profiles in quarters with a history of mastitis. Front. Microbiol. 7, 480 (2016).

15. Bhatt, V. D. et al. Milk microbiome signatures of subclinical mastitisu analysed by shotgun sequencing. J. Appl. Microbiol. 112(4), 639–650 (2012).

16. Hoque, M. N., Das, Z. C., Rahman, A. N. M. A., Haider, M. G., & Islam, M. A. Molecular characterization of *Staphylococcus aureus* strains in bovine mastitis milk in Bangladesh. Int. J. Vet. Sci. Med. 6(1), 53–60 (2018).

17. Jamali, H. et al. Invited review: Incidence, risk factors, and effects of clinical mastitis recurrence in dairy cows. J. Dairy Sci. 101(6), 4729–4746 (2018).

18. Swinkels, J. M., Lam, T. J. G. M., Green, M. J. & Bradley, A. J. Effect of extended cefquinome treatment on clinical persistence or recurrence of environmental clinical mastitis. Vet. J. 197, 682–687 (2013).

19. Hoque, M. N. et al. Different screening tests and milk somatic cell count for the prevalence of subclinical bovine mastitis in Bangladesh. Trop. Anim. Health Prod. 47(1), 79–86 (2015).

20. Patel, S. H. et al. Culture independent assessment of human milk microbial community in lactational mastitis. Sci. Rep. 7(1), 7804 (2017).

21. Egilmez, H. I. et al. Temperature-dependent virus lifecycle choices may reveal and predict facets of the biology of opportunistic pathogenic bacteria. Sci. Rep. 8(1), 1–13 (2018).

22. Lloyd, M. M. & Pespeni M. H. Microbiome shifts with onset and progression of Sea Star Wasting Disease revealed through time course sampling. Sci. Rep. 8(1), 16476 (2018).

23. Harvell, C. D. et al. Climate warming and disease risks for terrestrial and marine biota. Science 296, 2158–2162 (2002).

24. Gomes, F., Maria, J. S. & Mariana, H. Bovine mastitis disease/pathogenicity: evidence of the potential role of microbial biofilms. Pathogens Dis. 74(3), (2016).

25. Andrews, T., Neherm, D. A., Weicht, T. R. & Barlow, J. W. Mammary microbiome of lactating organic dairy cows varies by time, tissue site, and infection status. PLoS One 14(11), e0225001 (2019).

26. Seth, S., Välimäki, N., Kaski, S. & Honkela, A. Exploration and retrieval of whole-metagenome sequencing samples. Bioinformatics 30(17), 2471–2479 (2014).

27. Oniciuc, E. et al. The present and future of Whole Genome Sequencing (WGS) and Whole Metagenome Sequencing (WMS) for surveillance of antimicrobial resistant microorganisms and antimicrobial resistance genes across the food chain. Genes 9(5), 268 (2018).

28. Hong, C. et al. PathoScope 2.0: a complete computational framework for strain identification in environmental or clinical sequencing samples. Microbiome 2(1), 33 (2014).

29. Glass, E. M., Wilkening, J., Wilke, A., Antonopoulos, D. & Meyer, F. Using the metagenomics RAST server (MG-RAST) for analyzing shotgun metagenomes. Cold Spring Harb Protoc. 2010(1), 5368 (2010).

30. Liu, B., Zheng, D., Jin, Q., Chen, L. & Yang, J. VFDB 2019: a comparative pathogenomic platform with an interactive web interface. Nucleic Acids Res. 47(D1), D687–D692 ((2019).

31. Doster, E. et al. ResFinder 2.0: a database for classification of antimicrobial drug, biocide and metal resistance determinants in metagenomic sequence data. Nucleic Acids Res. 48(D1), D561–D569 (2019).

32. Kanehisa, M. et al. New approach for understanding genome variations in KEGG. Nucleic Acids Res. 47, D590–D595 (2019).

33. Rizzatti, G. et al. Proteobacteria: a common factor in human diseases. BioMed Res. Intl. 9351507, 7 (2017).

34. Boix-Amoros, A., Collado, M. C., Miram A. Relationship between Milk Microbiota, Bacterial Load, Macronutrients, and Human Cells during Lactation. Front. Microbiol. 7, 492 (2016).

35. Hoque, M. N. et al. Molecular characterization of *Staphylococcus aureus* strains in bovine mastitis milk in Bangladesh. Int. J. Vet. Sci. Med. 6, 53–60 (2018).

36. Segata N. On the Road to Strain-Resolved Comparative Metagenomics. mSystems 3(2), e00190–17 (2018).

37. Round, J. L. & Mazmanian, S. K. The gut microbiota shapes intestinal immune responses during health and disease. Nat. Rev. Immunol. 9, 313–323 (2009).

38. Nayfach, S. et al. New insights from uncultivated genomes of the global human gut microbiome. Nature 568, 505–510 (2019).

39. Mohr, E. L., Berhane, A., Zora, J. G. & Suchdev, P. S. *Acinetobacter baumannii* neonatal mastitis: a case report. J. Med. Case Rep. 8(1), 318 (2014).

40. Catozzi, C. et al. The microbiota of water buffalo milk during mastitis. PLoS One 12(9), e0184710 (2017).

41. Lee C. R. et al. “Biology of *Acinetobacter baumannii*: Pathogenesis, Antibiotic Resistance Mechanisms, and Prospective Treatment Options.” Front Cell Infect Microbiol. 13, 7(55) (2017).

42. Klaas, I. C. & Zadoks, R. N. An update on environmental mastitis: Challenging perceptions. Transbound. Emerging Dis. 65, 166 –185 (2018).

43. Asai, N. et al. *Pantoea dispersa* bacteremia in an immunocompetent patient: a case report and review of the literature. J. Med. Case Rep. 13, 33 (2019).

44. Ma, C. et al. Cow-to-mouse fecal transplantations suggest intestinal microbiome as one cause of mastitis. Microbiome 6(1), 200 (2018).

45. Maga, E. A., Weimer, B. C. & Murray, J. D. Dissecting the role of milk components on gut microbiota composition. Gut Microbes 4(2), 136–9 (2013).

46. Lurie-Weinberger & Gophna, M. N. Archaea in and on the human body: health implications and future directions. PLoS Pathog. 11(6), e1004833 (2015).

47. Ruiz, L., García-Carral, C. & Rodriguez, J. M. Unfolding the human milk microbiome landscape in the omics era. Front. Microbiol. 10, (2019).

48. Legault, B.A., Lopez-Lopez, A., Alba-Casado, J.C. et al. Environmental genomics of “*Haloquadratum walsbyi*” in a saltern crystallizer indicates a large pool of accessory genes in an otherwise coherent species. BMC Genomics 7, 171 (2006).

49. Addis, M. F., Tanca, A., Uzzau, S., Oikonomou, G., Bicalho, R. C. & Moroni, P. The bovine milk microbiota: insights and perspectives from -omics studies. Mol. Biosyst. 12, 2359–2372 (2016).

50. Fleitas, M. O. et al. Recent Advances in Anti-virulence Therapeutic Strategies with a Focus on Dismantling Bacterial Membrane Microdomains, Toxin Neutralization, Quorum-Sensing Interference and Biofilm Inhibition. Front. Cellular Infect. Microbiol. 9, 74 (2019).

51. Zeng, M. Y., Inohara, N. & Nunez, G. Mechanisms of inflammation-driven bacterial dysbiosis in the gut. Mucosal Immunol. 10(1), 18–26 (2017).

52. Peng, Y. et al. Roles of Hcp family proteins in the pathogenesis of the porcine extraintestinal pathogenic *Escherichia coli* type VI secretion system. Sci. Rep. 6, 26816 (2016).

53. Heurlier, K., Dénervaud, V., Pessi, G., Reimmann, C. & Haas, D. Negative control of quorum sensing by RpoN (σ54) in *Pseudomonas aeruginosa* PAO1. J. Bact. 185(7), 2227–2235 (2003).

54. Addis, M. F., Tanca, A., Uzzau, S., Oikonomou, G., Bicalho, R. C. & Moroni, P. The bovine milk microbiota: insights and perspectives from -omics studies. Mol. Biosyst. 12, 2359–2372 (2016).

55. Kamada, N., Chen, G. Y., Inohara, N. & Núñez, G. Control of pathogens and pathobionts by the gut microbiota. Nat. Immunol. 14, 685–690 (2013).

56. Fitzpatrick, D. & Walsh, F. Antibiotic resistance genes across a wide variety of metagenomes. FEMS Microbiol. Ecol. 92(2), fiv168 (2016).

57. Nobrega, D. B. et al. Prevalence and Genetic Basis of Antimicrobial Resistance in Non-aureus *Staphylococci* Isolated from Canadian Dairy Herds. Front. Microbiol. 9, 256 (2018).

58. Card, R., Zhang, J., Das, P., Cook, C., Woodford, N. & Anjum, M. F. Evaluation of an expanded microarray for detecting antibiotic resistance genes in a broad range of gram-negative bacterial pathogens. Antimicrob. Agents Chemother. 57, 458–465 (2013).

59. Van Hoek, A. H., Mevius, D., Guerra, B., Mullany, P., Roberts, A. P. & Aarts, H. J. Acquired antibiotic resistance genes: an overview. Front. Microbiol. 2, 203 (2011).

60. Turnbaugh, P. J. et al. A core gut microbiome in obese and lean twins. Nature 457(7228), 480–487 (2009).

61. Li, N. et al. Variation in raw milk microbiota throughout 12 months and the impact of weather conditions. Sci. Rep. 8(1), 2371 (2018).

62. Li, X., Ding, X. Z., Wan, Y. L., Liu, Y. M. & Du, G. Z. Comparative proteomic changes of differentially expressed whey proteins in clinical mastitis and healthy yak cows. Genet. Mol. Res. 13(3), 6593–6601 (2014).

63. Harris, S. & Levine, A. The p53 pathway: positive and negative feedback loops. Oncogene 24, 2899–2908 (2005).

64. Binnenkade, L., Lassak, J. & Thormann, K. M. Analysis of the BarA/UvrY two-component system in Shewanella oneidensis MR-1. PloS One, 6(9), e23440 (2011).

65. Argov, T. et al. Coordination of cohabiting phage elements supports bacteria–phage cooperation. Nat. Commun. 10, 5288 (2019).

66. Kelly, R.A. et al. Modelling changes in glutathione homeostasis as a function of quinone redox metabolism. Sci. Rep. 9, 6333 (2019).

67. Bar-Or, D., Bar-Or, R., Rael, L. T. & Brody, E. N. Oxidative stress in severe acute illness. Redox Biol. 4, 340–345 (2015).

68. Saleh, A., Macia Ortega, A. & Muotri, A. R. Transposable Elements, Inflammation and Neurological Disease. Front. Neurology 10, 894 (2019).

69. Head, S. R. et al. Library construction for next-generation sequencing: overviews and challenges. Biotechniques 56(2), 61–77 (2014).

70. Langmead, B. & Salzberg, S. L. Fast gapped-read alignment with Bowtie 2. Nat. Methods. 9, 357–359 (2012).

71. Francis, O. E. et al. Pathoscope: species identification and strain attribution with unassembled sequencing data. Genome Res. 23(10), 1721–1729 (2013).

72. Koh, H. An adaptive microbiome α-diversity-based association analysis method. Sci. Rep. 8(1), 18026 (2018).

73. Beck, J., Holloway, J. D. & Schwanghart, W. Under sampling and the measurement of beta diversity. Methods Ecol. Evol. 4(4), 370–382 (2013).

74. McMurdie, P. J. & Susan, H. Phyloseq: an R package for reproducible interactive analysis and graphics of microbiome census data. PLoS One 8(4), e61217 (2013).

75. Hu, Y. et al. OmicCircos: a simple-to-use R package for the circular visualization of multidimensional omics data. Cancer Informatics 13, 3–20 (2014).

