## Supplementary figures and images for "Microbiome Dynamics of Bovine Mastitis Progression and Genomic Determinants"

### Supplementary Figure 1

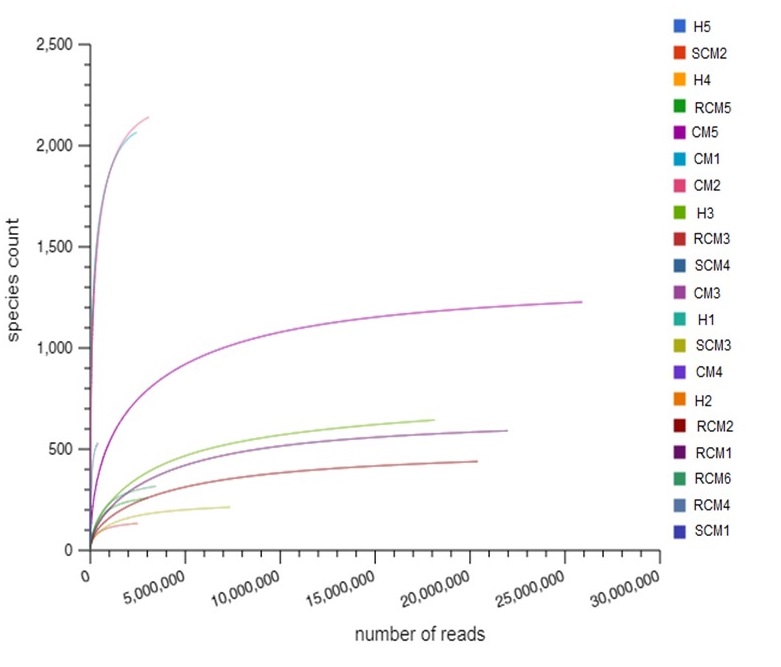

### Supplementary Figure 3

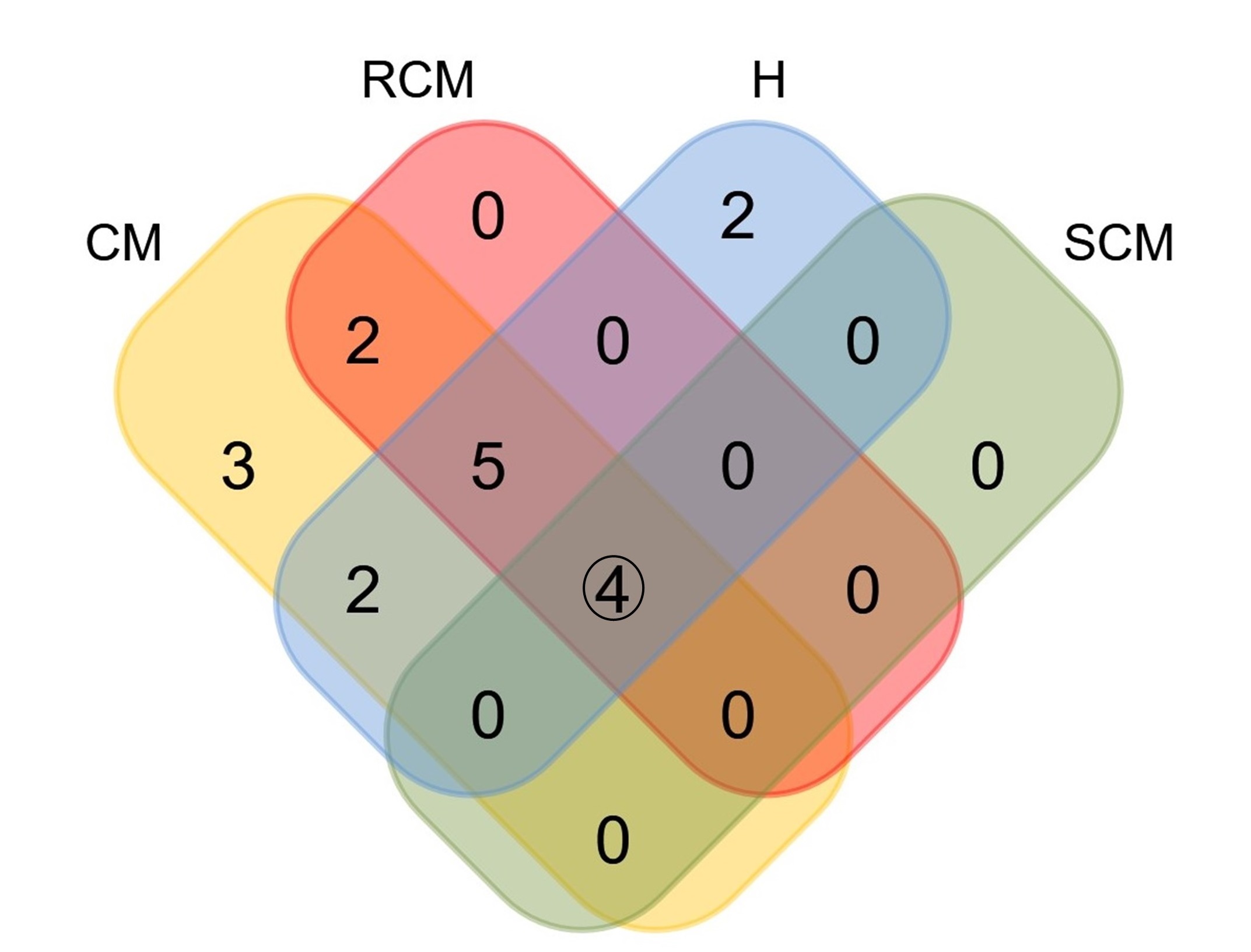

### Supplementary Figure 3

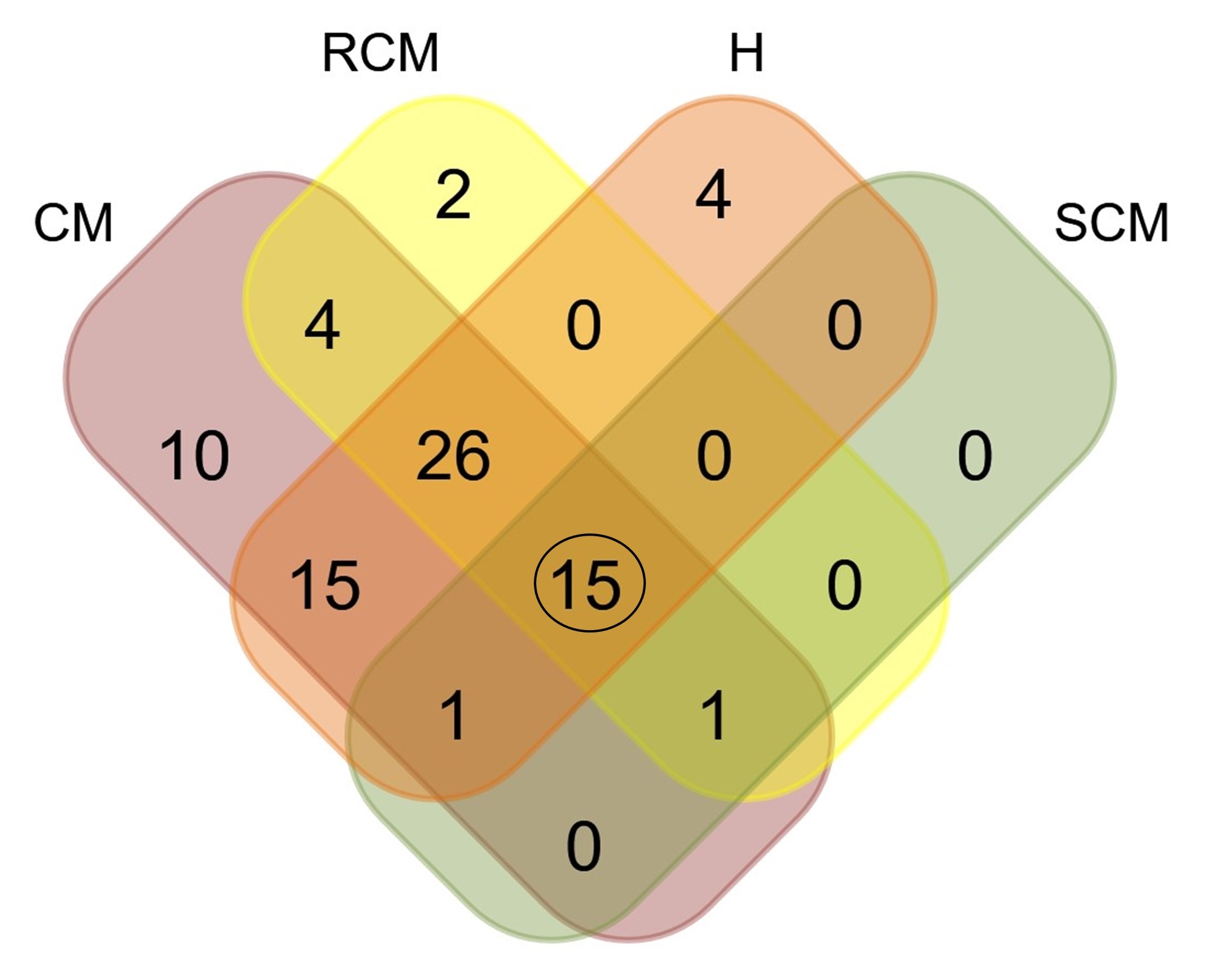

### Supplementary Figure 4

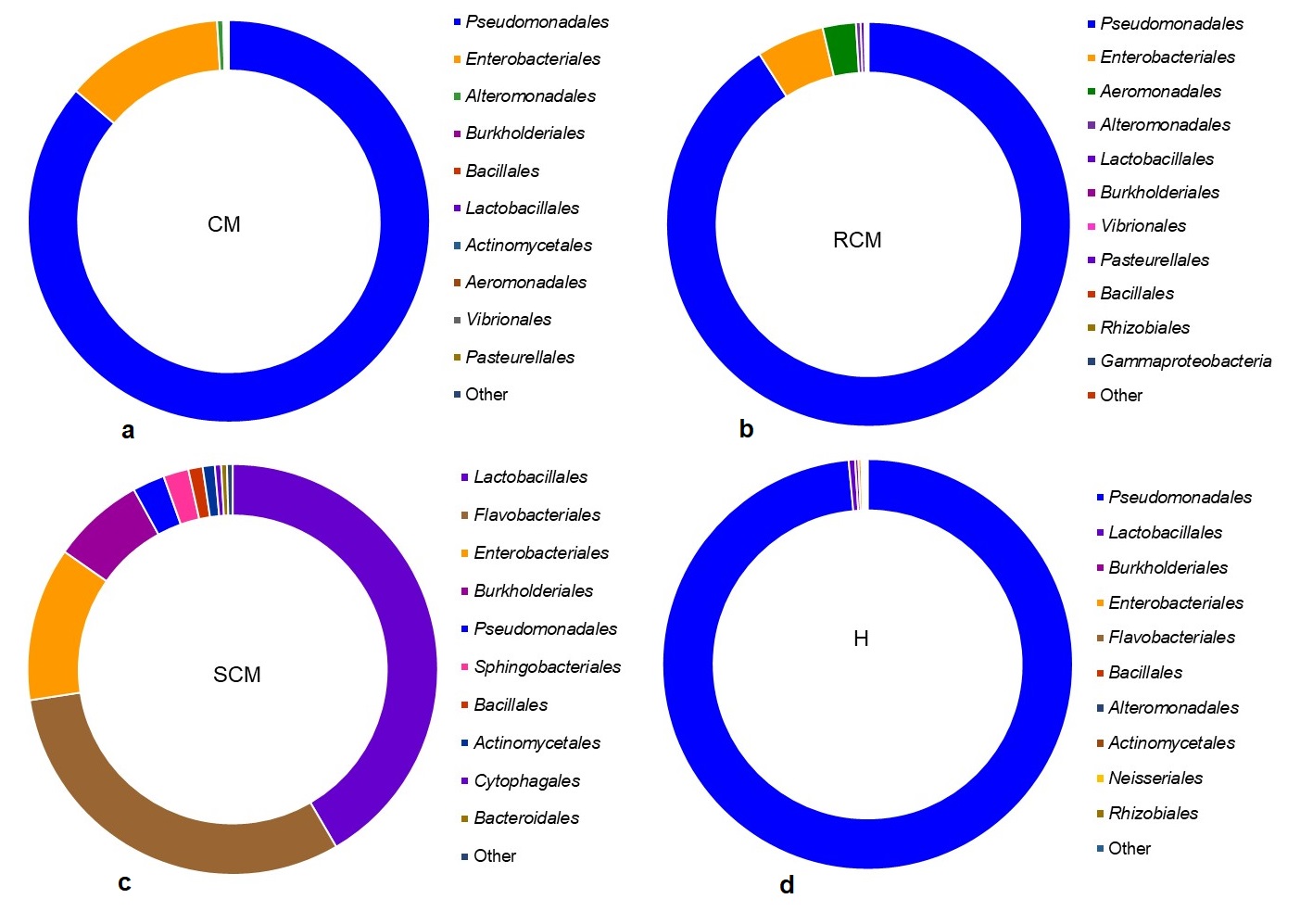
