## Supplementary Table 1 for "Microbiome Dynamics of Bovine Mastitis Progression and Genomic Determinants"

**Supplementary Table 1.** A total of 20 lactating crossbred cows (including 5 clinical mastitis, 6 recurrent clinical mastitis, 4 subclinical mastitis and 5 healthy) were selected for the study from five different districts of Bangladesh. The cow, farm, milk sample and reads obtained from whole metagenome sequencing (WMS) of each sample related information are given in the table below. Here, CM, Clinical mastitis; RCM, recurrent clinical mastitis; SCM, subclinical mastitis; H, Healthy milk; SW, Sahiwal crossbred; RCC, Red Chattogram Cattle; LZ, Local zebu; XHF, Holstein Friesian crossbred.

| **Cow ID** | **Sample ID** | **Farm Location** | **Farm ID** | **GIS (Longi-/Latitude)** | **No. of sample/farm** | **Breeds** | **Lactation (Days after calving)** | **Parity** | **Reads/sample (Before QC)** | **Reads/sample (after QC)** |
| --- | --- | --- | --- | --- | --- | --- | --- | --- | --- | --- |
| Cow 1 | CM1 | Gazipur | 1 | 24.19 N, 90.47 E | 1 | SW | 17 | 2 | 16784920 | 15532924 |
| Cow 2 | CM2 | Gazipur | 2 | 24.19 N, 90.47 E | 1 | RCC | 34 | 3 | 18607098 | 17157162 |
| Cow 3 | CM3 | Gazipur | 3 | 24.19 N, 90.47 E | 1 | LZ | 25 | 1 | 14769340 | 13810876 |
| Cow 4 | CM4 | Gazipur | 4 | 24.19 N, 90.47 E | 1 | XHF | 17 | 2 | 20090020 | 18727286 |
| Cow 5 | CM5 | Manikgonj | 5 | 23.86° N, 90.00° E | 1 | XHF | 9 | 4 | 39754738 | 37356232 |
| Cow 6 | RCM1 | Gazipur | 6 | 24.19 N, 90.47 E | 1 | SW | 48 | 2 | 28275998 | 26336406 |
| Cow 7 | RCM2 | Gazipur | 7 | 24.19 N, 90.47 E | 1 | RCC | 42 | 2 | 22676110 | 20364244 |
| Cow 8 | RCM3 | Chattogram | 8 | 22.20 N, 91.98 E | 1 | RCC | 15 | 3 | 17311874 | 15788554 |
| Cow 9 | RCM4 | Dhaka | 9 | 23.81 N, 90.41 E | 1 | XHF | 10 | 3 | 18497524 | 17110308 |
| Cow 10 | RCM5 | Manikgonj | 10 | 23.86° N, 90.00° E | 1 | SW | 32 | 1 | 18339188 | 16846638 |
| Cow 11 | RCM6 | Sirajgonj | 11 | 24.31° N, 89.57° E | 1 | XHF | 25 | 2 | 19507424 | 18088740 |
| Cow 12 | SCM1 | Chattogram | 12 | 22.20 N, 91.98 E | 1 | SW | 31 | 2 | 23563460 | 21888704 |
| Cow 13 | SCM2 | Dhaka | 13 | 23.81 N, 90.41 E | 1 | XHF | 13 | 5 | 19999578 | 18620308 |
| Cow 14 | SCM3 | Gazipur | 14 | 24.09 N, 90.42 E | 1 | SW | 41 | 3 | 2728026 | 2500796 |
| Cow 15 | SCM4 | Manikgonj | 15 | 23.86° N, 90.00° E | 1 | RCC | 22 | 2 | 34215394 | 31875744 |
| Cow 16 | H1 | Chattogram | 16 | 22.20 N, 91.98 E | 1 | XHF | 7 | 5 | 28491010 | 26161394 |
| Cow 17 | H2 | Chattogram | 17 | 22.34 N, 91.87 E | 1 | RCC | 32 | 3 | 19692182 | 18189320 |
| Cow 18 | H3 | Sirajgonj | 18 | 24.31° N, 89.57° E | 1 | XHF | 28 | 4 | 23857586 | 22114902 |
| Cow 19 | H4 | Dhaka | 19 | 23.81 N, 90.41 E | 1 | SW | 12 | 2 | 6772790 | 6009520 |
| Cow 20 | H5 | Gazipur | 20 | 24.19 N, 90.47 E | 1 | XHF | 23 | 3 | 22710894 | 21050232 |
