## Supplementary Table 2 for "Microbiome Dynamics of Bovine Mastitis Progression and Genomic Determinants"

**Supplementary Table 2:** Opportunistic and/or unreported strains in bovine mastitis milk metagenomes.

| **SL. No.** | **Opportunistic strains** | **Relative abundance** |
| --- | --- | --- |
| **Clinical mastitis (CM) milk** | | |
| 1 | Aeromonas veronii B565 | 19.16 |
| 2 | Pantoea dispersa EGD-AAK13 | 18.65 |
| 3 | Klebsiella oxytoca KA-2 | 11.7 |
| 4 | Kluyvera ascorbata ATCC 33433 | 7.27 |
| 5 | Aeromonas hydrophila YL17 | 6.54 |
| 6 | Kluyvera intermedia E86 | 4.53 |
| 7 | Aeromonas hydrophila subsp. hydrophila ATCC 7966 | 3.32 |
| 8 | Klebsiella aerogenes KCTC 2190 | 3.09 |
| 9 | Aeromonas hydrophila SSU | 2.81 |
| 10 | Aeromonas eucrenophila | 2.39 |
| 11 | Shewanella oneidensis MR-1 | 2.08 |
| 12 | Kluyvera cryocrescens NBRC 102467 | 1.67 |
| 13 | Enterobacter cloacae subsp. cloacae ATCC 13047 | 1.06 |
| 14 | Plautia stali symbiont | 0.93 |
| 15 | Citrobacter freundii CFNIH1 | 0.86 |
| 16 | Acinetobacter gyllenbergii NIPH 230 | 0.72 |
| 17 | gamma proteobacterium L18 | 0.71 |
| 18 | Serratia marcescens FGI94 | 0.68 |
| 19 | Kosakonia sacchari SP1 | 0.62 |
| 20 | Serratia marcescens subsp. marcescens Db11 | 0.59 |
| 21 | Pseudomonas alcaligenes OT 69 | 0.56 |
| 22 | Aeromonas salmonicida subsp. salmonicida A449 | 0.56 |
| 23 | Enterobacter lignolyticus SCF1 | 0.56 |
| **Recurrent clinical mastitis (RCM) milk** | | |
| 1 | Nocardia pseudobrasiliensis | 48.06 |
| 2 | Serratia marcescens subsp. marcescens Db11 | 16.22 |
| 3 | Nocardia mikamii NBRC 108933 | 15.19 |
| 4 | Corynebacterium bovis DSM 20582 = CIP 54.80 | 4.66 |
| 5 | Aeromonas hydrophila SSU | 3.11 |
| 6 | Aeromonas veronii B565 | 1.81 |
| 7 | Aeromonas hydrophila YL17 | 1.73 |
| 8 | Streptococcus salivarius | 1.64 |
| 9 | Bradyrhizobium japonicum 22 | 1.64 |
| 10 | Cupriavidus metallidurans CH34 | 1.55 |
| 11 | Glutamicibacter arilaitensis Re117 | 1.04 |
| 12 | Methylobacterium radiotolerans JCM 2831 | 0.78 |
| 13 | Nocardia veterana NBRC 100344 | 0.69 |
| **Subclinical mastitis (SCM) milk** | | |
| 1 | Chryseobacterium sp. Leaf405 | 19.06 |
| 2 | Chryseobacterium sp. CF299 | 13.68 |
| 3 | Chryseobacterium haifense DSM 19056 | 13.19 |
| 4 | Chryseobacterium greenlandense | 13.04 |
| 5 | Chryseobacterium sp. YR460 | 12.56 |
| 6 | Serratia marcescens subsp. marcescens Db11 | 8.3 |
| 7 | Citrobacter freundii CFNIH1 | 6.65 |
| 8 | Sediminibacterium salmoneum NBRC 103935 | 2.35 |
| 9 | Elizabethkingia anophelis NUHP1 | 2.2 |
| 10 | Streptococcus salivarius | 1.49 |
| 11 | Chryseobacterium koreense CCUG 49689 | 1.27 |
| 12 | Sphingobacterium spiritivorum ATCC 33861 | 1.23 |
| 13 | Empedobacter brevis NBRC 14943 = ATCC 43319 | 1.12 |
| 14 | Methyloglobulus morosus KoM1 | 0.75 |
| 15 | Lactobacillus algidus DSM 15638 | 0.75 |
