## Supplementary Table 3 for "Microbiome Dynamics of Bovine Mastitis Progression and Genomic Determinants"

| Sl. No | **CM milk metagenome** | | | | |
| --- | --- | --- | --- | --- | --- |
|  | **VFGs** | **Associated Bacteria** | **Taxon-Abund.** | **VFGs-Abund.** | **Correlation** |
| 1 | *ABZJ* | *Acinetobacter* | 68.59 | 9.65 | Non-parametric Kruskal Wallis Test,  p = 0.001 |
| 2 | *pvd*L | *Pseudomonas* | 17.69 | 5.11 |  |
| 3 | *chp*A | *Pseudomonas* | 17.69 | 3.17 |  |
| 4 | *mot*B*/omp*A | *Acinetobacter*, *Salmonella*,  *Yersinia, Vibrio*, *Aeromonas* | 69.26 | 2.11 |  |
| 5 | *pilJ* | *Pseudomonas* | 17.69 | 1.83 |  |
| 6 | *flh*A | *Pseudomonas*, *Yersinia*, *Aeromonas, Bacillus* | 17.83 | 1.67 |  |
| 7 | *gac*S | *Pseudomonas* | 17.69 | 1.56 |  |
| 8 | *icm*F1 | *Pseudomonas,* *Burkholderia* | 17.70 | 1.51 |  |
| 9 | *fli*F | *Yersinia, Vibrio*, , *Campylobacter* | 0.038 | 1.40 |  |
| 10 | *alg8* | *Pseudomonas* | 17.69 | 1.39 |  |
| **RCM milk metagenome** | | | | | |
| 1 | *ABZJ* | *Acinetobacter* | 72.61 | 8.81 | Non-parametric Kruskal Wallis Test,  p = 0.001 |
| 2 | *pvdL* | *Pseudomonas* | 18.21 | 5.28 |  |
| 3 | *chpA* | *Pseudomonas* | 18.21 | 2.91 |  |
| 4 | *motB/ompA* | *Acinetobacter*, *Aeromonas, Salmonella*,  *Yersinia, Vibrio* | 75.90 | 2.34 |  |
| 5 | *pilJ* | *Pseudomonas* | 18.21 | 1.81 |  |
| 6 | *flhA* | *Pseudomonas*, *Yersinia*, *Aeromonas, Bacillus* | 20.94 | 1.78 |  |
| 7 | *icmF1* | *Pseudomonas,* *Burkholderia* | 18.22 | 1.52 |  |
| 8 | *gacS* | *Pseudomonas* | 18.21 | 1.45 |  |
| 9 | *fleQ* | *Pseudomonas, Aeromonas Escherichia* | 21.52 | 1.32 |  |
| 10 | *alg8* | *Pseudomonas* | 18.21 | 1.31 |  |
| **SCM milk metagenome** | | | | | |
| 1 | *EcSMS35* | *Escherichia* | 3.978 | 5.28 | Non-parametric Kruskal Wallis Test,  p = 0.001 |
| 2 | *cpsB* | *Klebsiella,* *Streptococcus, Enterococcus* | 2.49 | 2.61 |  |
| 3 | *stgC* | *Escherichia, Salmonella* | 4.57 | 2.10 |  |
| 4 | *upaG/ehaG* | *Escherichia* | 3.978 | 2.11 |  |
| 5 | *iroC* | *Escherichia, Klebsiella, Salmonella, Yersinia* | 5.56 | 2.08 |  |
| 6 | *Rfb* | *Streptococcus* | 1.13 | 1.78 |  |
| 7 | *iroN* | *Escherichia, Klebsiella, Streptococcus, Shigella* | 6.02 | 1.70 |  |
| 8 | *elf* | *Escherichia* | 3.978 | 1.60 |  |
| 9 | *hcp* | *Escherichia* | 3.978 | 1.48 |  |
| 10 | *cfaC* | *Escherichia* | 3.978 | 1.41 |  |
| **H milk metagenome** | | | | | |
| 1 | *ABZJ* | *Acinetobacter* | 73.66 | 16.92 | Non-parametric Kruskal Wallis Test,  p = 0.001 |
| 2 | *hcp2* | *Pseudomonas* | 24.01 | 7.65 |  |
| 3 | *hsiC2* | *Pseudomonas* | 24.01 | 7.61 |  |
| 4 | *rpoN* | *Pseudomonas,*  *Dichelobacter* | 24.02 | 4.87 |  |
| 5 | *algG* | *Pseudomonas* | 24.01 | 4.28 |  |
| 6 | *alg8* | *Pseudomonas* | 24.01 | 3.93 |  |
| 7 | *motB/ompA* | *Acinetobacter*, *Salmonella*, *Vibrio*, *Yersinia*, *Aeromonas* | 73.72 | 3.43 |  |
| 8 | *oprF* | *Pseudomonas* | 24.01 | 3.33 |  |
| 9 | *fleQ* | *Pseudomonas, Escherichia, Aeromonas* | 24.04 | 3.31 |  |
| 10 | *fliM* | *Salmonella, Aeromonas, Vibrio, Yersinia* | 0.062 | 3.10 |  |

**Microbiome Dynamics of Bovine Mastitis Progression and Genomic Determinants**

M. Nazmul Hoque, Arif Istiaq, M. Shaminur Rahman, M. Rafiul Islam, Azraf Anwar, AMAM Zonaed Siddiki, Munawar Sultana, Keith A. Crandall, M. Anwar Hossain

**Supplementary Table 3**: Correlation between taxonomic abundance and associated VFGs in four metagenomes (Top 10 abundant genes in each metagenome).
