## Supplementary material for "Microbiome Dynamics of Bovine Mastitis Progression and Genomic Determinants": Main Figure Legends

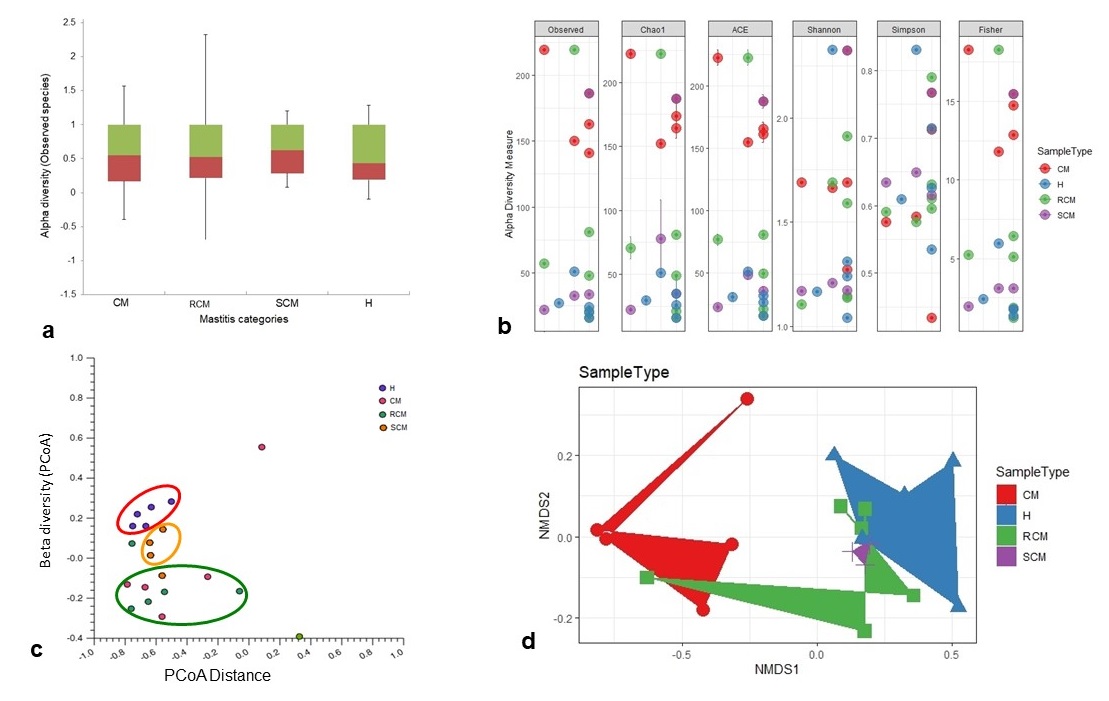

**Fig. 1. Differences in microbiome community between bovine mastitis and healthy milk.** (a) Box plots showing significant (P_Observed_ = 0.032) differences in observed species and/or strains richness in clinical mastitis (CM), recurrent clinical mastitis (RCM), subclinical mastitis (SCM) and healthy (H) milk samples. (b) Alpha diversity measured using the observed species, Chao 1, ACE, Shannon, Simpson and Fisher diversity indices through PathoScope (PS) analysis revealed distinct microbiome diversity across the samples of four metagenomes (p = 0.003, Kruskal-Wallis test). (c) Principal coordinates analysis (PCoA) measured on the Bray-Curtis distance method using MG-RAST (MR) tool (genus-level) colored samples by populations. Each dot represents an individual, and colors indicate the populations in four metagenomes. (d) Non-metric multidimensional scaling (NMDS) ordination plots showing the clear separation between CM, RCM, SCM and H milk samples as measured by weighted-UniFrac distance on PS data at strain level. In both cases, the CM microbiomes clustered more closely to RCM microbiomes (green circle in c, and red and green colored zones in d) followed by SCM microbiomes (red circle in c and violet zone in d), and however, more distantly clustered with H milk microbes (pink circle in c, and blue zone in d). Statistical analysis using Kruskal–Wallis tests showed significant microbial diversity variations across the four metagenomes (p = 0.001, Kruskal Wallis test).

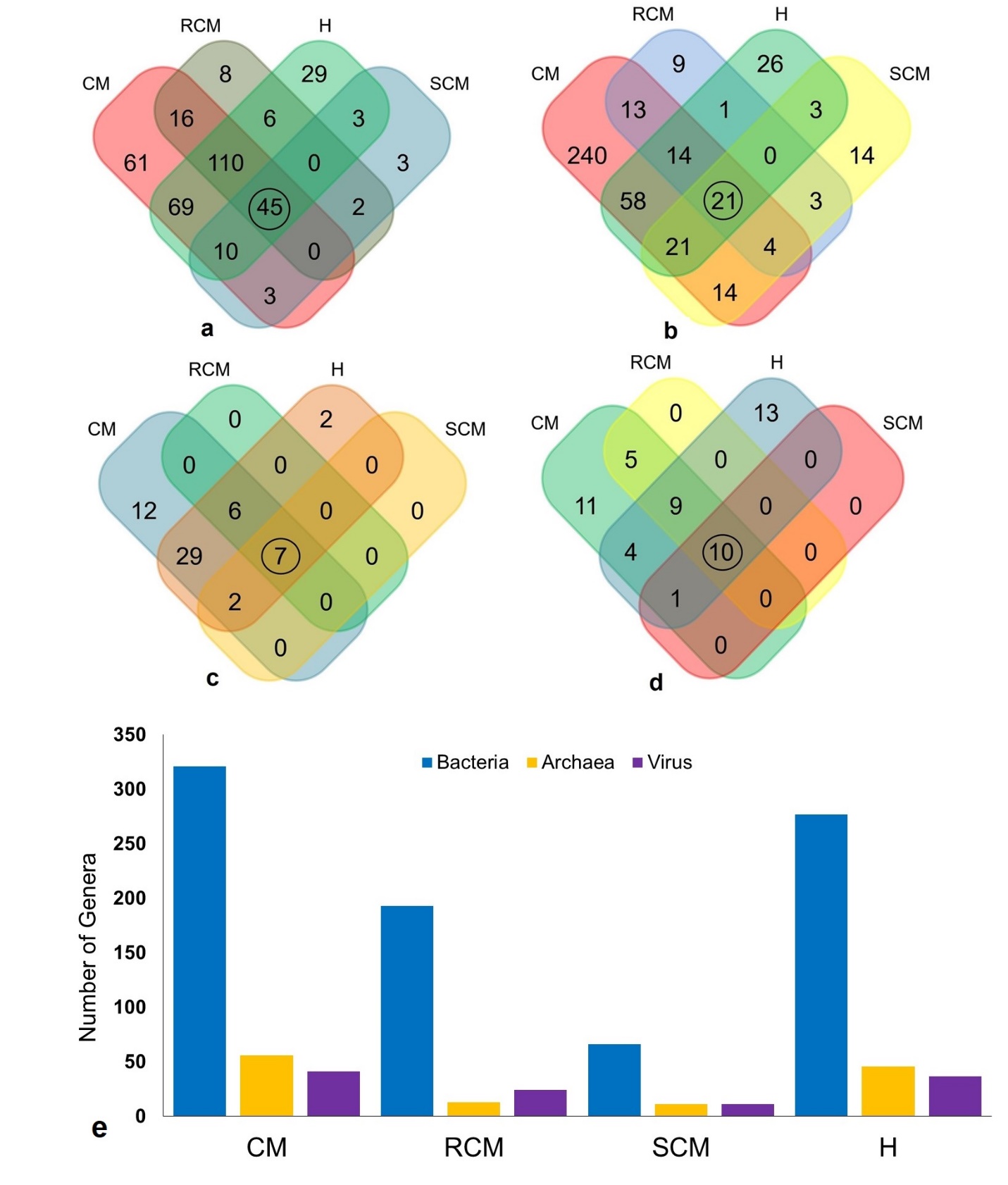
**Fig. 2. Taxonomic composition of bovine mastitis and healthy milk microbiome.** Venn diagrams representing the core unique and shared microbiomes in CM, RCM, SCM and H milk samples. (a) Venn diagram showing unique and shared bacterial genera by MR analysis, (b) Venn diagram comparison of bacteria at strain level as measured through PS analysis, (c & d) Venn diagrams representing unique and shared archaeal and viral genera, respectively found in mastitis (CM, RCM, SCM) and H milk samples as analyzed with MR pipeline. Microbiome sharing between the conditions are indicated by black circles. (e) Dynamic changes in the composition of bacteria, archaea and viruses in four metagenomes at genus level. Each bar plot represents total genera of bacteria, archaea and viruses detected in the respective metagenome.

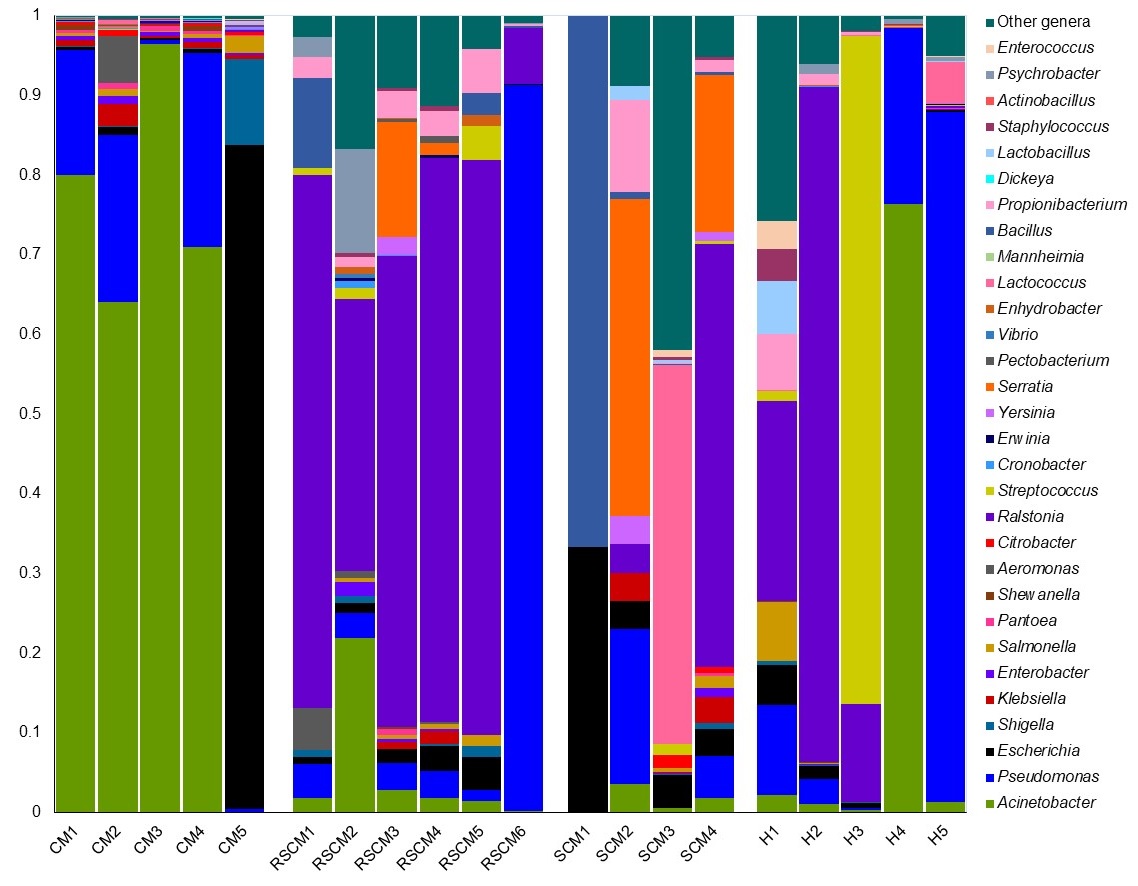

**Fig. 3**. **The genus level taxonomic profile bacteria in bovine mastitis and healthy milk.** Stacked bar plots showing the relative abundance and distribution of the 30 most abundant genera, with ranks ordered from bottom to top by their increasing proportion among the CM, RCM, SCM and H milk samples. Only the 29 most abundant genera are shown in the legend, with the remaining genera grouped as ‘Other genera’. Each stacked bar plot represents the abundance of bacteria in each sample of the corresponding category. The distribution and relative abundance of the bacterial genera in four metagenomes are also available in Supplementary Data 1.

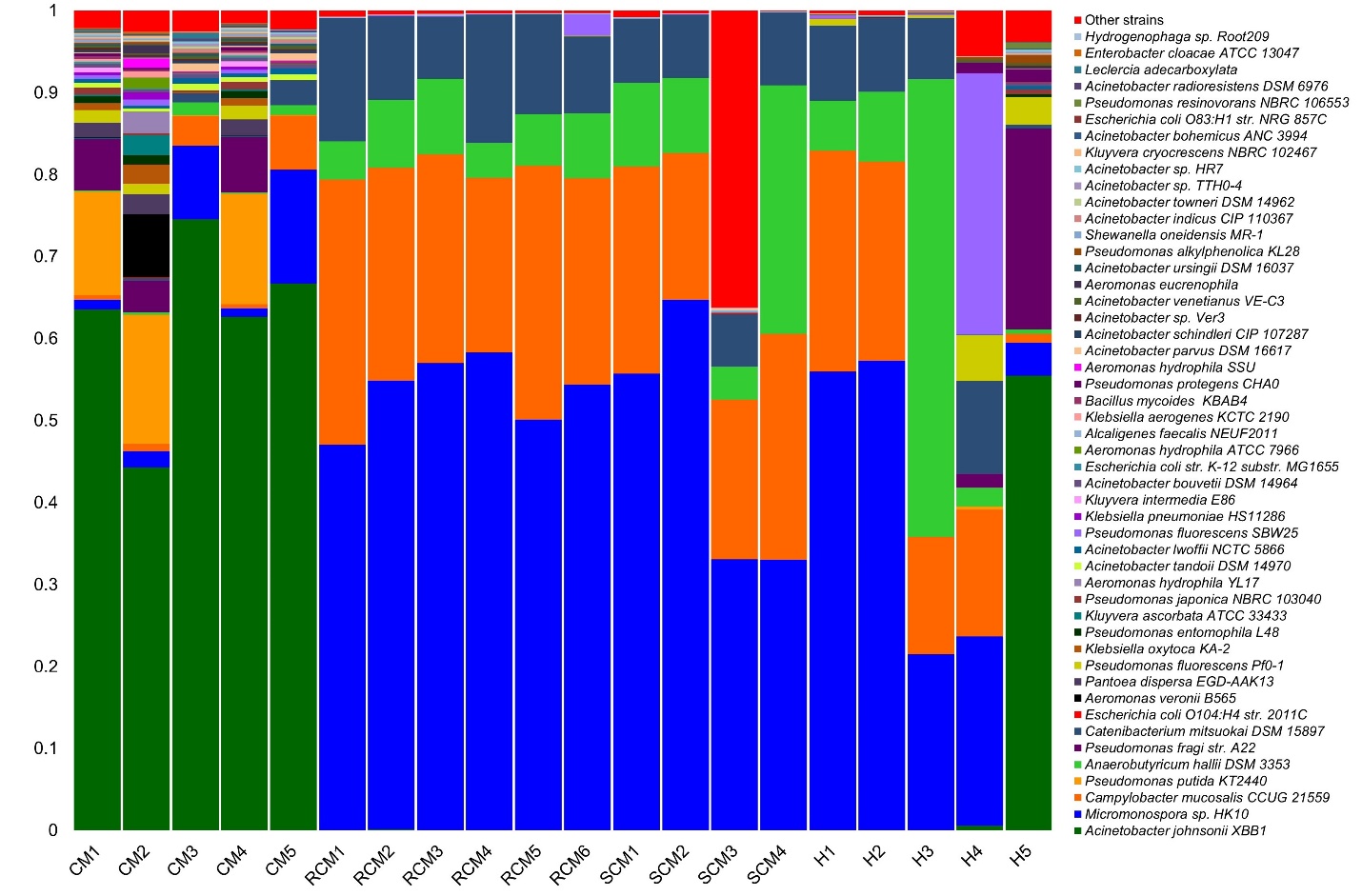

**Fig. 4**. **The strain level taxonomic profile bacteria in bovine mastitis and healthy milk.**

Stacked bar plots showing the relative abundance and distribution of the 50 most abundant strains, with ranks ordered from bottom to top by their increasing proportion among the CM, RCM, SCM and H milk samples. Only the 49 most abundant strains are shown in the legend, with the remaining strains grouped as ‘Other strains’. Each stacked bar plot represents the abundance of bacteria in each sample of the corresponding category. The distribution and relative abundance of the bacterial strains in four metagenomes are also available in Supplementary Data 1.

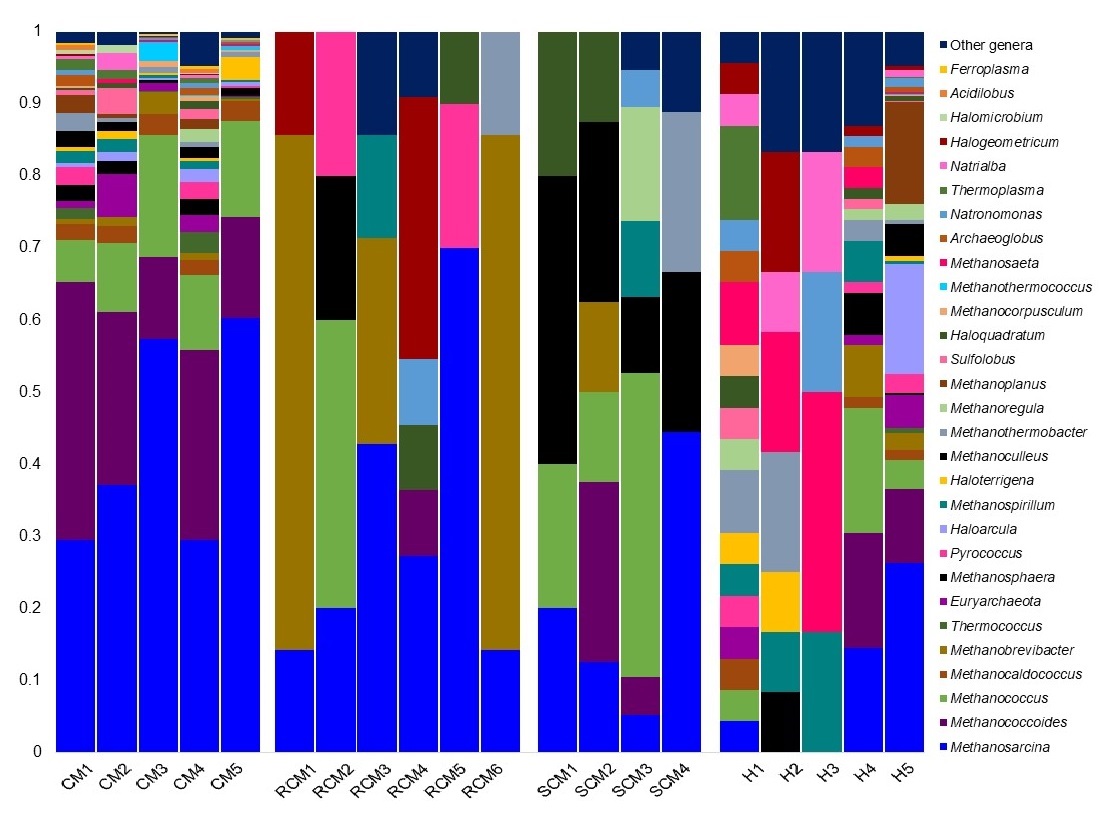

**Fig. 5**. **The taxonomic profile archaea in bovine mastitis and healthy milk.** Bar plots showing the distribution and relative abundance of top 30 most abundant archaeal genera, with ranks ordered from bottom to top by their increasing proportion among the CM, RCM, SCM and H milk samples. Only the 29 most abundant genera are shown in the legend, with the remaining strains grouped as ‘Other genera’. Each stacked bar plot represents the abundance of archaea in each sample of the corresponding category, and notable differences in the archaeal populations are those where the taxon is abundant in one sample category and effectively undetected in the other sample types. The distribution and relative abundance of the archaeal genera in four metagenomes are also available in Supplementary Data 1.

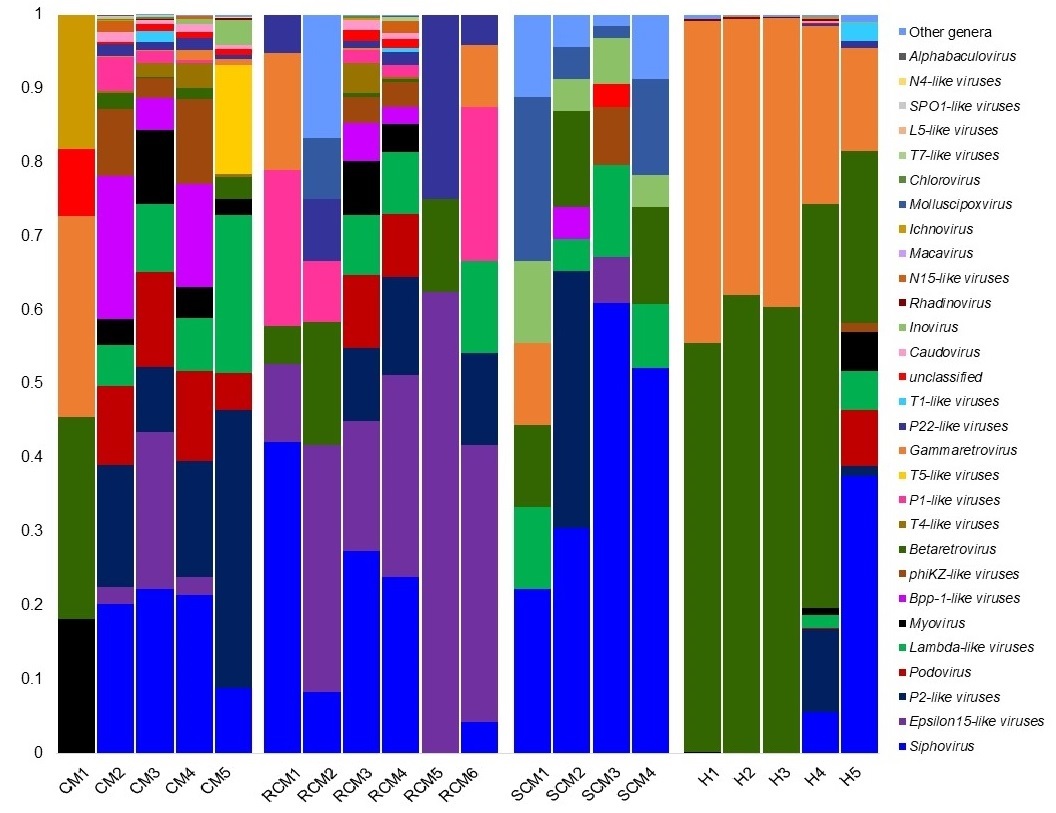

**Fig. 6**. **The taxonomic profile virus in bovine mastitis and healthy milk.** Taxonomic distribution of 30 viral genera detected in all of the 20 samples of CM, RCM, SCM and H milk metagenomes. The most abundant viral genera are sorted by descending order of the relative abundance. Each stacked bar plot represents the abundance of viruses in each sample of the corresponding category, and notable differences in the viral populations are those where the taxon is abundant in one sample category and effectively undetected in the other sample types. The distribution and relative abundance of the viral genera in four metagenomes are also available in Supplementary Data 1.

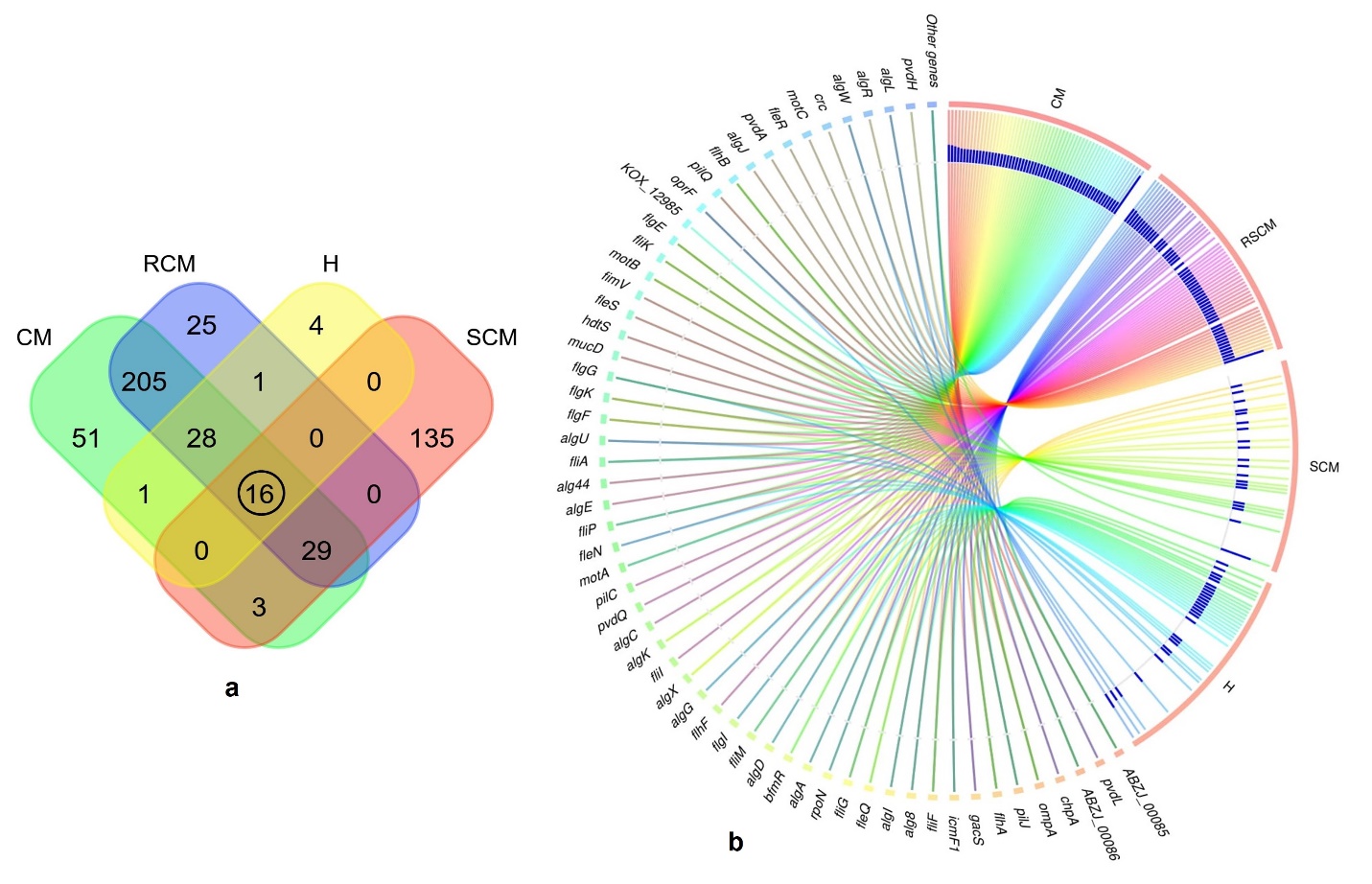

**Fig. 7.** **Virulence factors associated genes (VFGs) detected in bovine mastitis and healthy milk metagenomes.** Metagenome sequencing data was used to search for open reading frames (ORFs) compared against the VFDB database to identify the VFGs with over 95% sequence identity. (a) Venn diagram showing unique and shared VFGs detected in CM, RCM, SCM and H milk metagenomes. VFGs sharing between the conditions are indicated by black circle. (b) The circular plot illustrates the distribution of top abundant 50 VFGs found across the four metagenomes. VFGs in the respective metagenome group are represented by different colored ribbons, and the inner blue bars indicate their respective relative abundances. The CM associated microbiomes had the highest number of VFGs followed by RCM, SCM and H metagenomes. The distribution and relative abundance of the VFGs in four metagenomes are also available in Supplementary Data 2.

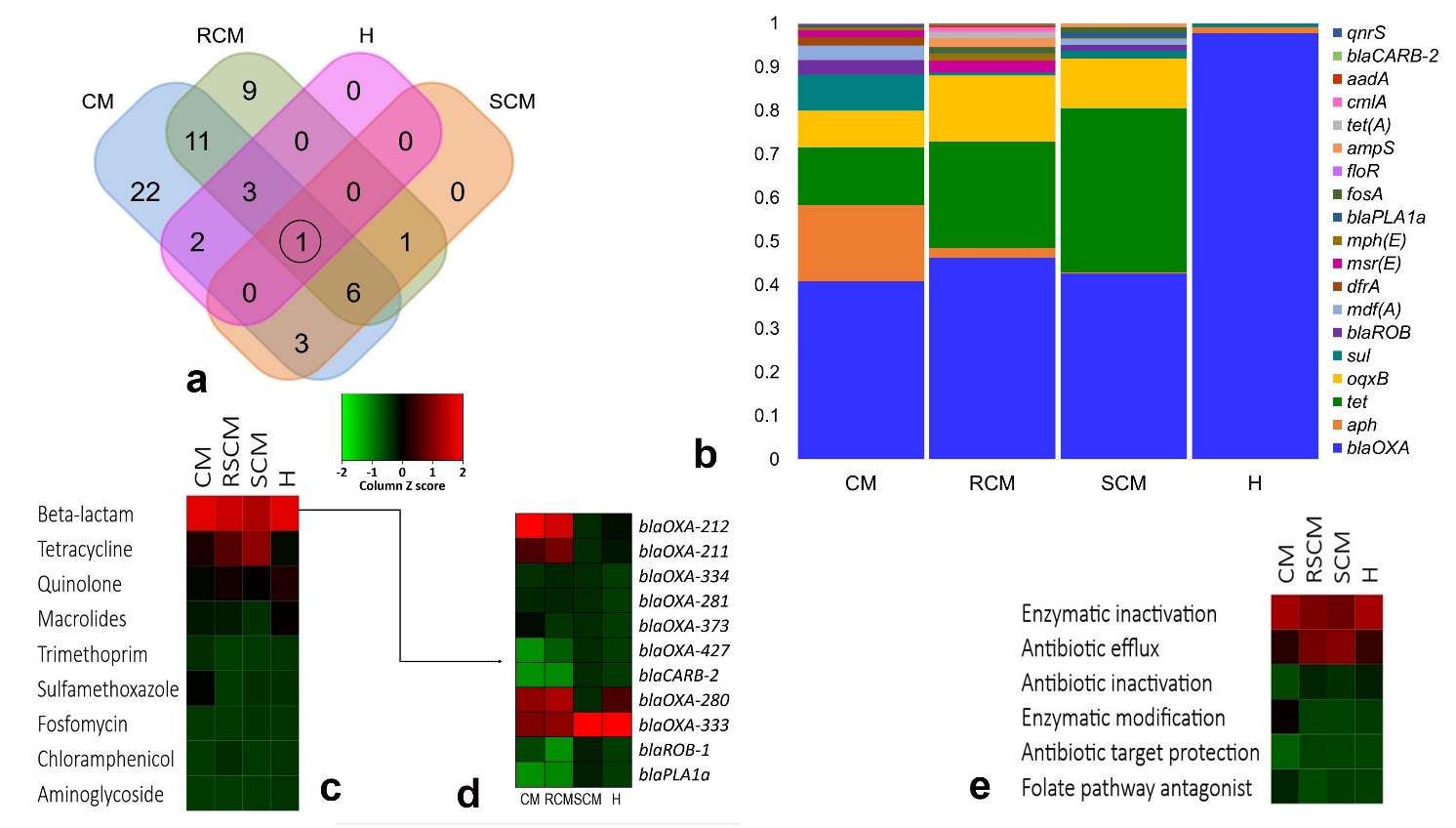

**Fig. 8.** **Antibiotic resistance genes (ARGs) in bovine mastitis and healthy milk metagenomes.** Metagenome sequencing data was used to search for open reading frames (ORFs) compared against the ResFinder database to identify ARGs with over 95% sequence identity. (a) Venn diagram showing unique and shared ARGs detected in CM, RCM, SCM and H milk metagenomes. ARGs sharing between the conditions are indicated by black circle. (b) Stacked bar plots showing the relative abundance and distribution of the ARGs across the four metagenomes. (c) Heatmap showing the classes of antibiotics associated with the identified ARGs in bovine mastitis and healthy milk microbiomes. (d) Different beta-lactam genes resistant genes found in four metagenomes. (e) The possible mechanisms of actions/pathways of antimicrobial resistance found in this metagenome sequences. The color code indicates the presence and completeness of each gene, expressed as a value (Z score) between -2 (low abundance), and 2 (high abundance). The red color indicates the highest abundance whilst light green cells accounts for lower abundance of the respective genes in each metagenome.

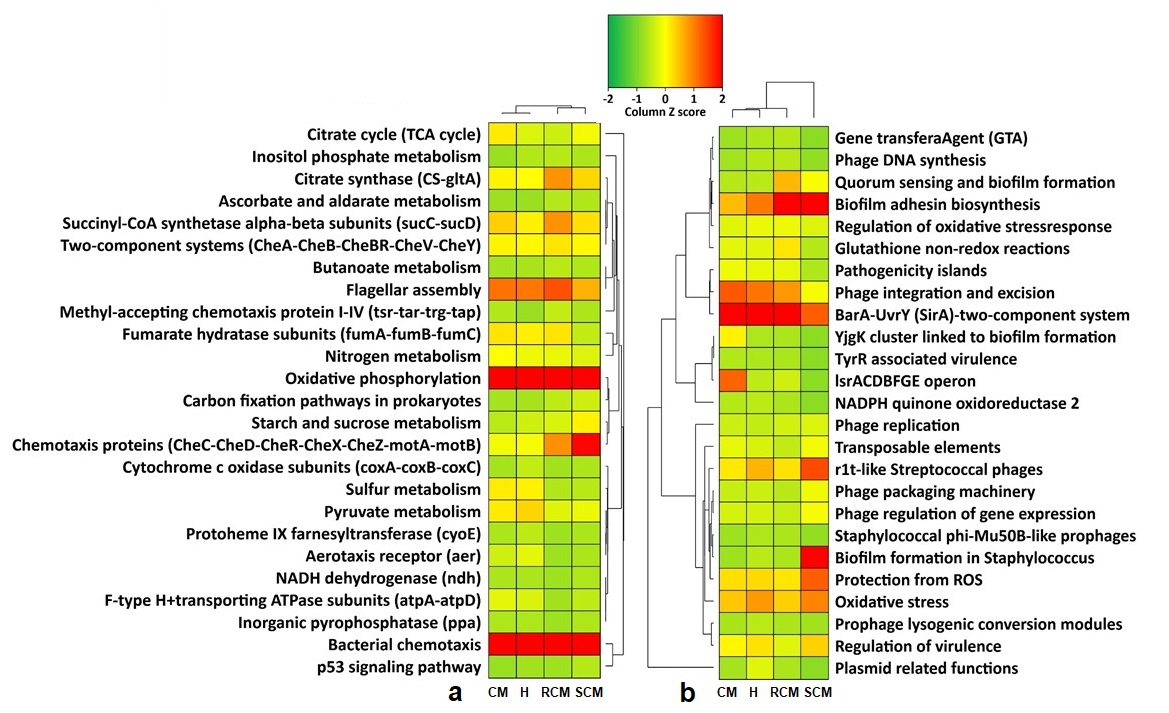

**Fig. 9: Projection of the mastitis and healthy milk metagenomes onto KEGG pathways and SEED subsystems.** The WMS reveals differences in metabolic functional pathways in CM, RCM, SCM and H milk microbiomes. (a). Heatmap showing the average relative abundance hierarchical clustering of the predicted KEGG Orthologs (KOs) functional pathways of the microbiota across all samples. (b). Comparison of metagenomic profiles of CM, RCM, SCM and H milk microbiotas at different levels of SEED subsystems (up to functions). The selected subsystems showing significant (p<0.05) differences among four sample groups is shown. The color bar at the top represents the relative abundance of putative genes. The color code indicates the presence and completeness of each KEGG and SEED module, expressed as a value (Z score) between -2 (low abundance), and 2 (high abundance). The red color indicates the highest abundant patterns of KOs and SEED subsystems, whilst light green cells accounts for less abundant KOs and SEED modules in that particular sample. More information on metabolic functional potentials (KOs and SEED modules) of the microbiomes in four metagenome groups are available in Supplementary Data 2.
