## Supplementary Figure Legends for "Microbiome Dynamics of Bovine Mastitis Progression and Genomic Determinants"

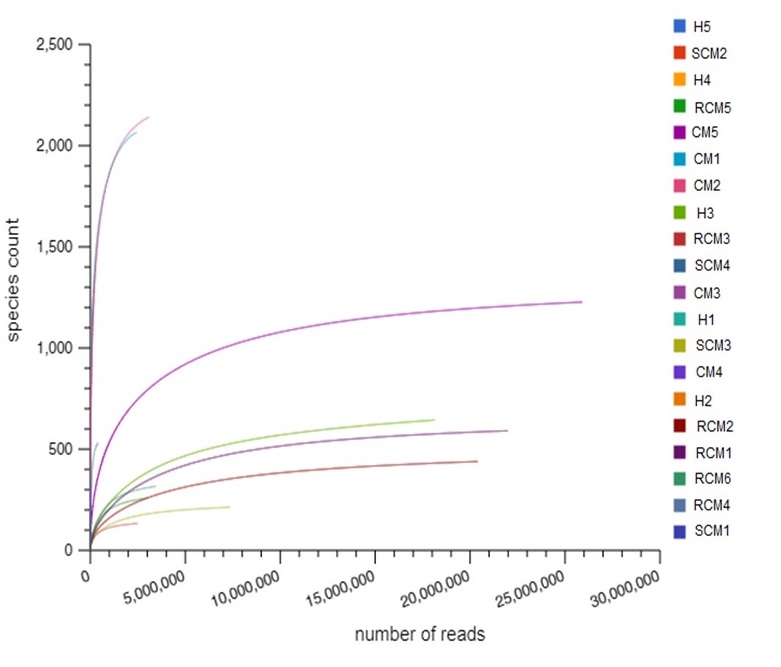

**Supplementary Figure 1.** Bovine mastitis and healthy milk microbiome diversity. Rarefaction curves showing the influence of sequencing depth (number of reads per sample, X axis) on species richness (Y axis) in clinical mastitis (CM), recurrent clinical mastitis (RCM), subclinical mastitis (SCM) and healthy (H) milk samples. The rarefaction curves representing the number of species per sample indicated that the sequencing depth was sufficient enough to fully capture the microbial diversity as existed.

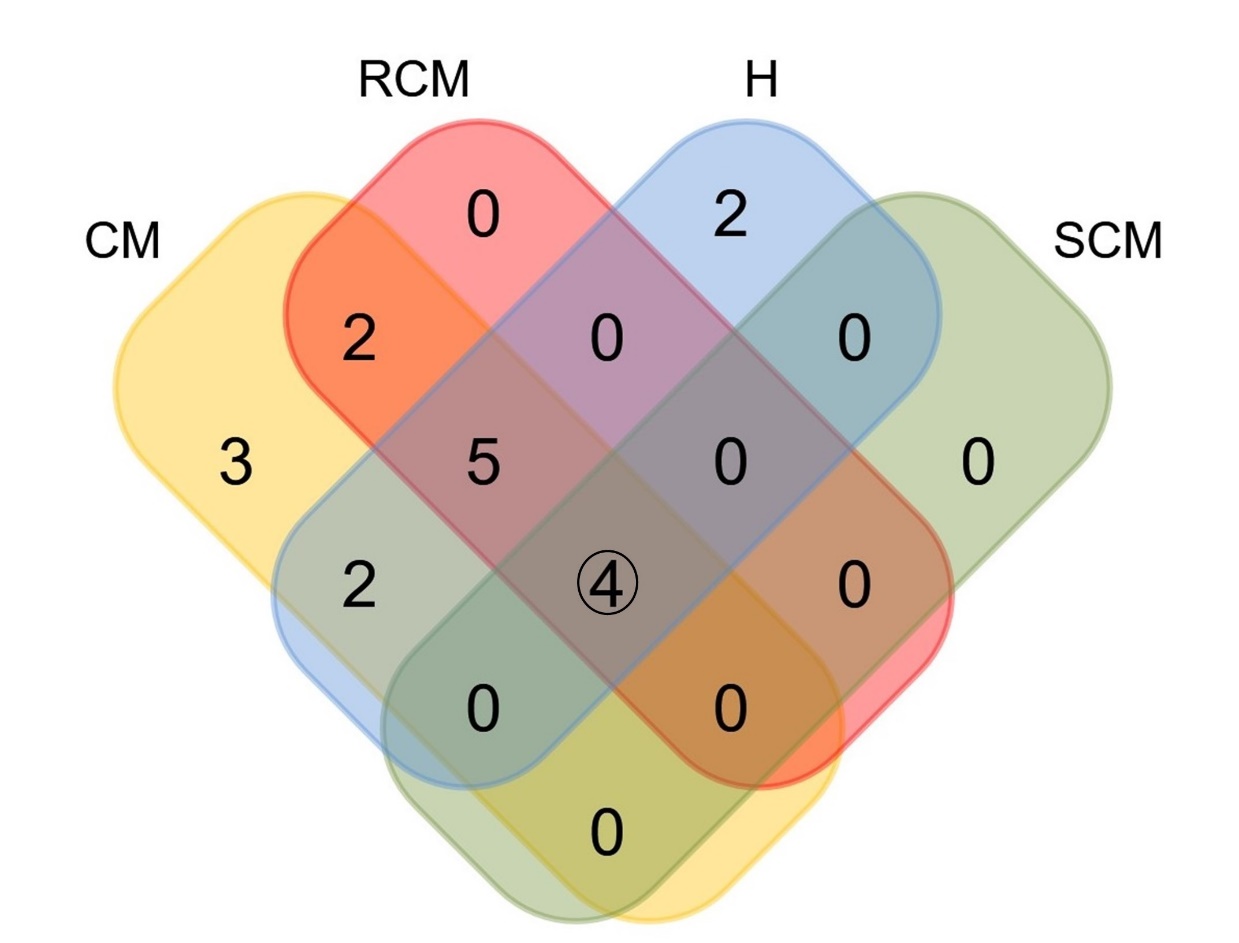

**Supplementary Figure 2. Taxonomic composition of bovine mastitis and healthy milk microbiome (at phylum level).** Venn diagram representing the core unique and shared bacterial phyla in CM, RCM, SCM and H milk samples. Microbiome sharing between the conditions are indicated by black circles.

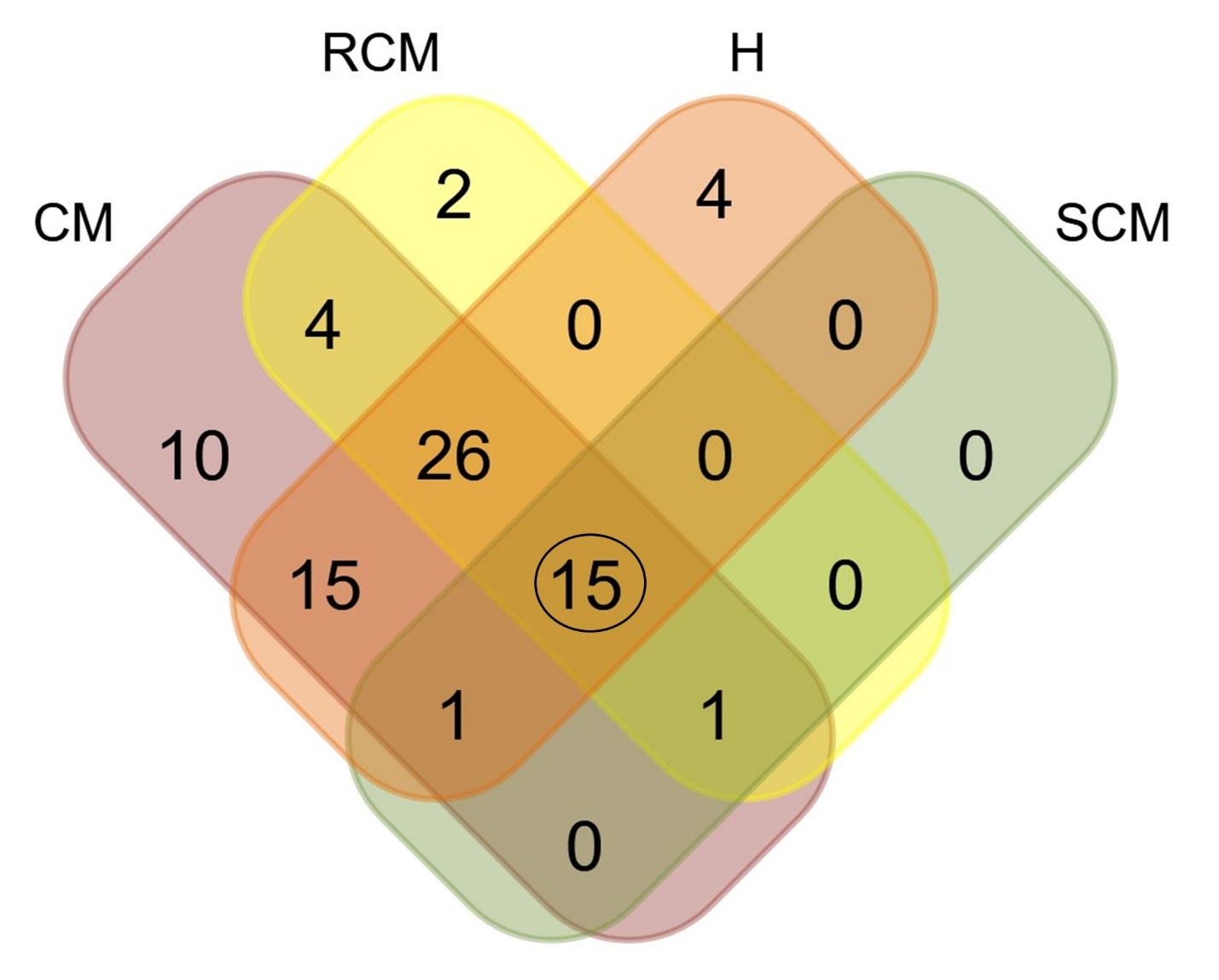

**Supplementary Figure 3**. **Taxonomic composition of bovine mastitis and healthy milk microbiome (at order level).** Venn diagram representing the core unique and shared bacterial orders in CM, RCM, SCM and H milk samples. Microbiome sharing between the conditions are indicated by black circles.

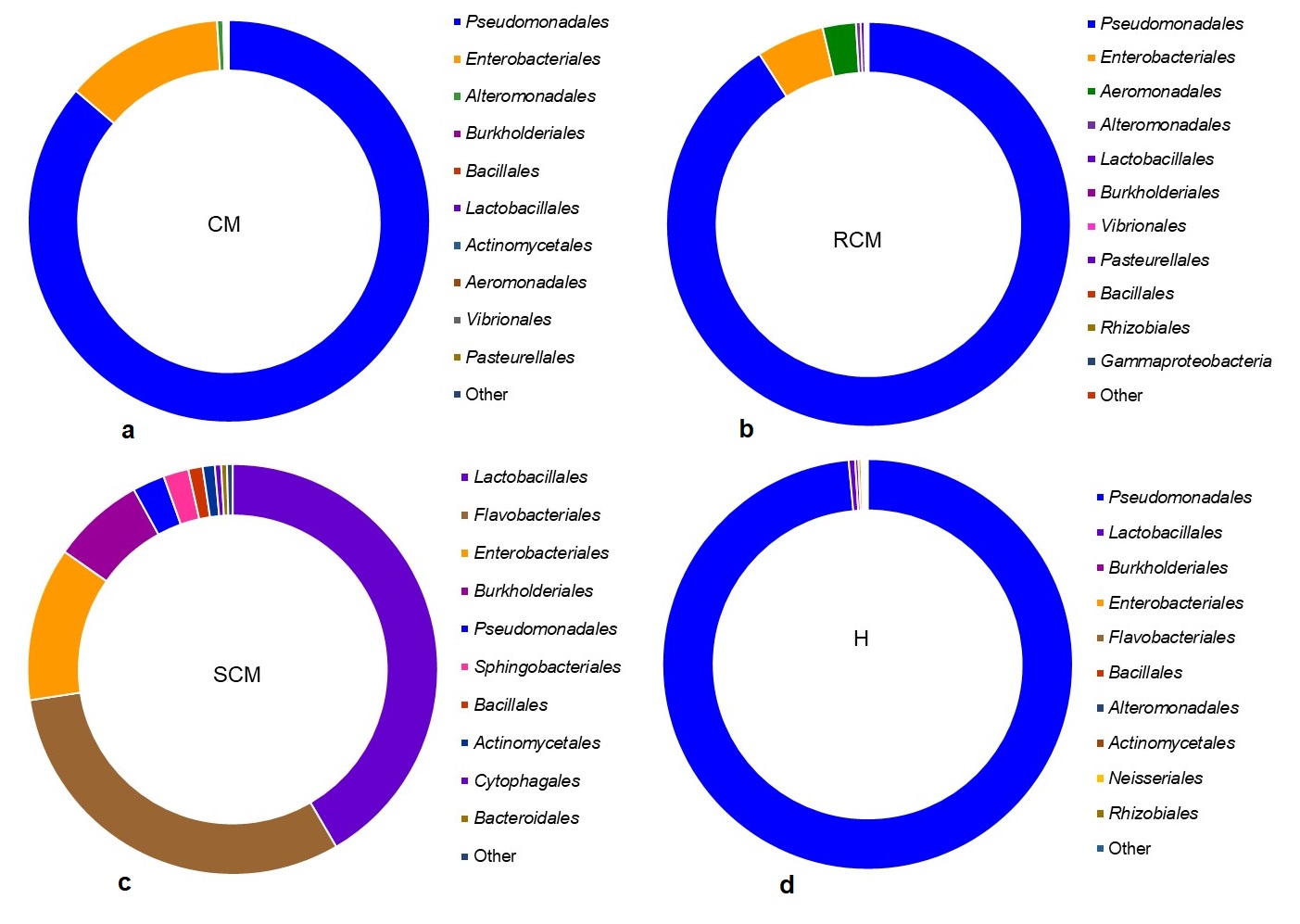

**Supplementary Figure 4**. **The taxonomic profile bacteria (at order level) in bovine mastitis and healthy milk.**  The circular plots showing to abundant bacterial orders in (a) clinical mastitis (CM), (b) recurrent clinical mastitis (RCM), (c) subclinical mastitis (SCM) and (d) healthy (H) milk samples. Different color codes indicate the relative abundance of the respective orders in all metagenome groups. The distribution and relative abundance of the bacterial orders in four metagenomes are also available in Supplementary Data 2.
